# The SWI/SNF chromatin remodeling assemblies BAF and PBAF differentially regulate cell cycle exit and cellular invasion *in vivo*

**DOI:** 10.1101/2021.03.01.433447

**Authors:** Jayson J. Smith, Yutong Xiao, Nithin Parsan, Taylor N. Medwig-Kinney, Michael A. Q. Martinez, Frances E. Q. Moore, Nicholas J. Palmisano, Abraham Q. Kohrman, Mana Chandhok Delos Reyes, Rebecca C. Adikes, Simeiyun Liu, Sydney A. Bracht, Wan Zhang, Kailong Wen, Paschalis Kratsios, David Q. Matus

## Abstract

Chromatin remodelers such as the SWI/SNF complex coordinate metazoan development through broad regulation of chromatin accessibility and transcription, ensuring normal cell cycle control and cellular differentiation in a lineage-specific and temporally restricted manner. Mutations in genes encoding the structural subunits of chromatin, such as histone subunits, and chromatin regulating factors (CRFs) are associated with a variety of disease mechanisms including cancer metastasis, in which cancer co-opts cellular invasion programs functioning in healthy cells during development. Here we utilize *Caenorhabditis elegans* anchor cell (AC) invasion as an *in vivo* model to identify the suite of chromatin agents and CRFs that promote cellular invasiveness. We demonstrate that the SWI/SNF ATP-dependent chromatin remodeling complex is a critical regulator of AC invasion, with pleiotropic effects on both G_0_ cell cycle arrest and activation of invasive machinery. Using targeted protein degradation and enhanced RNA interference (RNAi) vectors, we show that SWI/SNF contributes to AC invasion in a dose-dependent fashion, with lower levels of activity in the AC corresponding to aberrant cell cycle entry and increased loss of invasion. Our data specifically implicate the SWI/SNF BAF assembly in the regulation of the G_0_ cell cycle arrest in the AC, whereas the SWI/SNF PBAF assembly promotes AC invasion via cell cycle-independent mechanisms, including attachment to the basement membrane (BM) and activation of the pro-invasive *fos-1*/FOS gene. Together these findings demonstrate that the SWI/SNF complex is necessary for two essential components of AC invasion: arresting cell cycle progression and remodeling the BM. The work here provides valuable single-cell mechanistic insight into how the SWI/SNF assemblies differentially contribute to cellular invasion and how SWI/SNF subunit-specific disruptions may contribute to tumorigeneses and cancer metastasis.

**SUMMARY STATEMENT:** Cellular invasion through the basement membrane by the *C. elegans* anchor cell requires both BAF and PBAF SWI/SNF assemblies to arrest the cell cycle and promote the expression of pro-invasive genes.

## INTRODUCTION

Cellular invasion through basement membranes (BMs) is a critical step in metazoan development and is important for human health and fitness. Early in hominid development, trophoblasts must invade into the maternal endometrium for proper blastocyst implantation (1). In the context of immunity, leukocytes become invasive upon injury or infection to travel between the bloodstream and interstitial tissues (2,3). Atypical activation of invasive behavior is associated with a variety of diseases, including rheumatoid arthritis wherein fibroblast-like synoviocytes adopt invasive cellular behavior, leading to the expansion of arthritic damage to previously unaffected joints (4,5). Aberrant activation of cell invasion is also one of the hallmarks of cancer metastasis (6).

A variety of *in vitro* and *in vivo* models have been developed to study the process of cellular invasion at the genetic and cellular levels. *In vitro* invasion assays typically involve 3D hydrogel lattices, such as Matrigel, through which cultured metastatic cancer cells will invade in response to chemo-attractants (7). Recently, microfluidic systems have been integrated with collagen matrices to improve these *in vitro* investigations of cellular invasion (8). While *in vitro* invasion models provide an efficient means to study the mechanical aspects of cellular invasion, they are currently unable to replicate the complex microenvironment in which cells must invade during animal development and disease. A variety of *in vivo* invasion models have been studied, including cancer xenograft models in mouse (9–11) and zebrafish (12,13), each having their respective benefits and drawbacks. Over the past ∼15 years, *Caenorhabditis elegans* anchor cell (AC) invasion has emerged as a powerful alternative model due to its visually tractable single-cell nature (**Fig 1 A**) (14).

**Figure 1.**
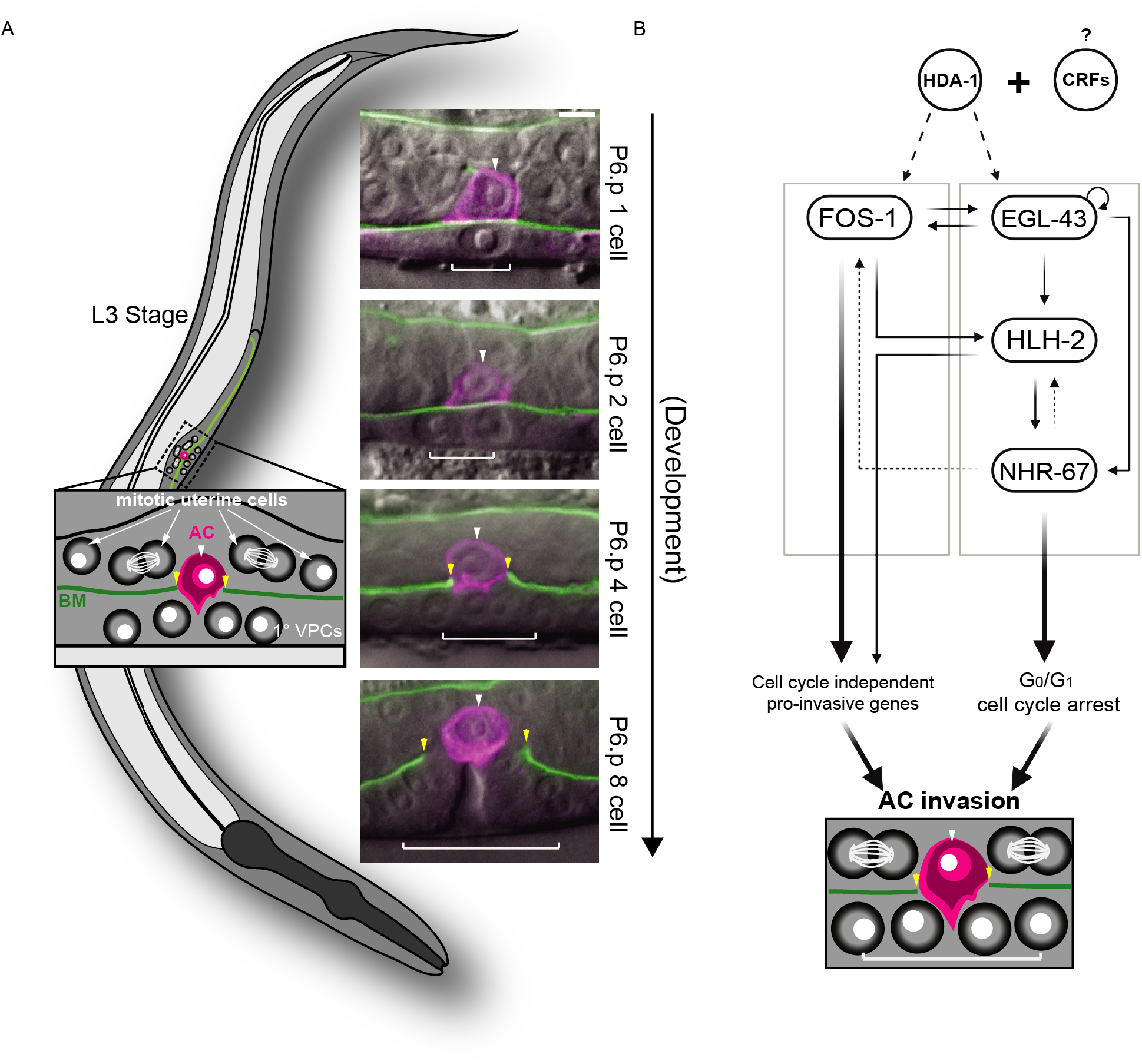
Summary of *C. elegans* AC invasion through the underlying BM and AC GRN. **(A)** Schematic depicting AC invasion in the mid-L3 stage of *C. elegans* development (left) and micrographs demonstrating the coordination of AC (magenta, *cdh-3p::PH::mCherry*) invasion through the BM (green, *laminin::GFP*) with primary vulval development. The fluorescent AC-specific membrane marker and BM marker are overlaid on DIC in each image. White arrowheads indicate AC(s), yellow arrowheads indicate boundaries of breach in BM, and white brackets indicate 1° VPCs. **(B)** Overview of the transcription factor GRN governing AC invasion (22,24), which consists of cell cycle-independent (*fos-1*) and dependent (*egl-43, hlh-2,* and *nhr-67*) subcircuits, which together with *hda-1* promote pro-invasive gene expression and maintain cell cycle arrest in the AC. In this and all subsequent figures, scale bar, 5 μm.

Previous work demonstrated a high degree of evolutionary conservation in the cell-autonomous mechanisms underlying BM invasion (3,15), including basolateral polarization of the F-actin cytoskeleton/cytoskeletal regulators and the expression of matrix metalloproteinases (MMPs) (16–21). Moreover, in order to breach the BM, the AC requires the expression of transcription factors (TFs), whose human homologs are common to metastatic cancers, including *egl-43* (EVI1/MEL), *fos-*1 (FOS), *hlh-2* (E/Daughterless), and *nhr-67* (TLX/Tailless) (22) (**Fig 1 B**). In addition to the expression of pro-invasive genes, there is increasing evidence that cells must also arrest in the cell cycle to adopt an invasive phenotype (23). Our previous work has demonstrated that the AC must terminally differentiate and arrest in the G_0_/G_1_ phase of the cell cycle to invade the BM and make contact with the underlying primary vulval precursor cells (1° VPCs) (22,24). The regulatory mechanisms that couple G_0_/G_1_ cell cycle arrest with the ability of a cell to invade the BM remain unclear.

Cell-extrinsic and cell-intrinsic factors, such as chromatin remodeling complexes and TFs, can influence the decision to maintain a plastic cell fate or to undergo cell fate specification and terminal differentiation. The determination of a cell to remain plastic or specify is in part the consequence of a complex, genome-wide antagonism between Polycomb group (PcG) transcriptional repression and Trithorax group (TrxG) transcriptional activation (25–27). For example, the binding of pioneer transcription factors OCT4 and SOX2 to target DNA in order to retain pluripotency in murine embryonic stem cells is the indirect consequence of the regulation of chromatin accessibility at these target regions (28). A recent study has shown that chromatin accessibility of enhancers in crucial cell cycle genes which promote the G_1_/S transition, including Cyclin E and E2F transcription factor 1, is developmentally restricted to reinforce terminal differentiation and cell cycle exit during *Drosophila melanogaster* pupal wing morphogenesis (29). In *C. elegans* myogenesis, the SWItching defective/Sucrose Non-Fermenting (SWI/SNF) ATP-dependent chromatin remodeling complex, a member of the TrxG family of complexes, both regulates the expression of the MyoD transcription factor (*hlh-1*) and acts redundantly to promote differentiation and G_0_ cell cycle arrest with core cell cycle regulators: cullin 1 (CUL1/*cul-1)*, cyclin-dependent kinase inhibitor 1 (*cki-1)*, FZR1 (*fzr-*1), and the RB transcriptional corepressor (RBL1/*lin-*35). The importance of regulation of chromatin states and the activity of CRFs for the acquisition and implementation of differentiated behaviors is also reflected in the *C. elegans* AC, as previous work has shown that the histone deacetylase *hda-1* (HDAC1/2) is required for pro-invasive gene expression and therefore differentiated cellular behavior and invasion (24) (**Fig 1 B**). Given these findings, a comprehensive investigation of the regulatory mechanism(s) governing AC invasion should include a thorough description of the suite of CRFs required for G_0_/G_1_ cell cycle arrest and invasive differentiation in the AC.

In this study we perform an RNA interference (RNAi) screen of *C. elegans* CRFs, specifically focusing on genes involved in chromatin structure and remodeling or histone modification. We identify 82 genes whose RNAi depletion phenotype resulted in a significant AC invasion defect. Among the CRFs identified as significant regulators of AC invasion, the SWI/SNF complex emerged as the most well-represented single chromatin remodeling complex. RNAi knockdown of subunits specific to the SWI/SNF core (*swsn-1* and *snfc-5/swsn-5*), and both BAF (BRG/BRM-Associated Factors; *swsn-8/let-526*) and PBAF (Polybromo Associated BAF; *pbrm-1* and *swsn-7*) assemblies resulted in penetrant loss of AC invasion. We generated fluorescent reporter knock-in alleles of subunits of the core (*GFP::swsn-4*) and BAF (*swsn-8::GFP*) assembly of the SWI/SNF complex using CRISPR/Cas9-mediated genome engineering. These alleles, used in conjunction with an endogenously labeled PBAF (*pbrm-1::eGFP*) assembly subunit, enabled us to determine the developmental dynamics of the SWI/SNF ATPase and assembly-specific subunits, gauge the efficiency of various SWI/SNF knockdown strategies, and asses inter- and intra-complex regulation. Using improved RNAi constructs and an anti-GFP nanobody degradation strategy (30), we demonstrated that the cell autonomous contribution of the SWI/SNF complex to AC invasion is dose dependent. This finding parallels similar studies in cancer (31–34) and *C. elegans* mesoblast development (35). Examination using a CDK activity sensor (36) revealed assembly-specific contributions to AC invasion: whereas BAF promotes AC invasion in a cell cycle-dependent manner, PBAF contributes to invasion in a cell cycle-independent manner. Finally, we utilized the auxin-inducible degron (AID) system combined with PBAF RNAi to achieve strong PBAF subunit depletion in the AC, which resulted in loss of both AC invasion and adhesion to the BM. Together, these findings provide insight into how the SWI/SNF complex assemblies may contribute to distinct aspects of proliferation and metastasis in human malignancies.

## RESULTS

### An RNAi screen of 269 CRFs identified SWI/SNF as a key regulator of AC invasion

To identify the suite of CRFs that, along with *hda-1,* contribute to AC invasion, we generated an RNAi sub-library of 269 RNAi clones from the complete Vidal RNAi library and a subset of the Ahringer RNAi library (37,38) targeting genes implicated in chromatin state, chromatin remodeling, or histone modification (**Table S1**) (**Fig 1 B**). Because CRFs act globally to control gene expression, we screened each RNAi clone by high-resolution differential interference contrast (DIC) and epifluorescence microscopy in a uterine-specific RNAi hypersensitive background containing labeled BM (*laminin::GFP*) and an AC reporter (*cdh-3p::PH::mCherry*) (**Fig 1 A; Table S1**) (14,22,24,39). This genetic background allowed us to limit the effect of transcriptional knockdown of CRFs to only affect the AC and the neighboring uterine tissue, and only for a time period following the specification of the AC (39). As the neighboring uterine cells do not contribute to the invasion program (14), AC invasion defects following RNAi treatments are indicative of cell autonomous pro-invasive gene function (24,39). In wild-type animals, by the time the 1° fated P6.p vulval precursor cell has divided twice (P6.p 4-cell stage), 100% of ACs have successfully breached the underlying BMs and made contact with the P6.p grand-daughters (14). While the AC also always invades in the uterine-specific RNAi hypersensitive strain we utilized for our CRF screen, there is a low penetrance delay, such that at the P6.p 4-cell stage, when we scored invasion, 2% (2/100 animals) still had an intact BM. Thus, we used this baseline defect as a statistical reference point for this genetic background and defined the cut-off threshold for significant defects in invasion from RNAi depletion of CRFs in our screen as those RNAi clones that resulted in a ∼13% or greater defect in invasion (4/30 animals, Fisher’s exact test = 0.0252). By this threshold, we recovered 82 CRFs (30.5% of total CRFs screened) that significantly regulate AC invasion, suggesting a general role for regulation of chromatin states in the acquisition of invasive behavior (**Table S2**). Interestingly, five subunits of the broadly conserved SWI/SNF chromatin remodeling complex were recovered as significant regulators of AC invasion: *swsn-1*(SMARCC1/SMARCC2; 23% AC invasion defect), *swsn-5/snfc-5* (SMARCB1; 20% AC invasion defect), *swsn-7* (ARID2; 23% AC invasion defect), *swsn-8/let-526* (ARID1A/ARID1B; 23% AC invasion defect), and *pbrm-1* (PBRM1; 20% AC invasion defect) (**Table S2**). As such, SWI/SNF is well-represented in our CRF screen, with representation of the core (*swsn-1* and *swsn-5*), BAF (*swsn-8*) and PBAF (*pbrm-1* and *swsn-7*) assemblies. Given the prevalence of SWI/SNF subunits recovered as significant regulators of AC invasion in our RNAi screen and the crucial role SWI/SNF plays in the regulation of animal development (40–45), tumorigenesis (32,46–48), and cell cycle control (35,49–52), we chose to focus our efforts on characterizing the role of the SWI/SNF complex in promoting AC invasion.

To confirm our RNAi results implicating the SWI/SNF complex in the promotion of AC invasion, we obtained two temperature sensitive hypomorphic alleles, *swsn-1(os22)* and *swsn-4(os13)* (41), and scored for defects in AC invasion in a genetic background containing both BM (*laminin::GFP*) and AC (*cdh-3p::mCherry::moeABD*) reporters. While we observed no defects in AC invasion in animals grown at the permissive temperature (15°C) (**Fig S1 A**), animals containing hypomorphic alleles for core subunits *swsn-1* and *swsn-4* cultured at the restrictive temperature (25°C) displayed defects in 20% (10/50) and 24% (12/50) of animals, respectively. These results with the *swsn-1(os22)* allele corroborated our *swsn-1(RNAi)* data from the CRF RNAi screen. Since neither of the RNAi libraries used to compose the CRF screen (see above) contained a *swsn-4(RNAi)* clone, results with the *swsn-4(os13)* allele also supplement data from our CRF RNAi screen by suggesting that AC invasion depends on the expression of the sole *C. elegans* SWI/SNF ATPase subunit in addition to the 5 subunits identified in the screen (**Fig S1 B**).

### Improved RNAi vectors revealed distinct contributions of SWI/SNF subunits to AC invasion

Though many SWI/SNF assemblies have been described in mammalian and other systems, including BAF, PBAF, esBAF, GBAF, nBAF, and npBAF) (53), to date, BAF and PBAF are the only SWI/SNF assemblies that have been described in *C. elegans*. Both assemblies consist of core subunits (SWSN-1, SWSN-4, SWSN-5) and accessory subunits (DPFF-1, SWSN-2.1/HAM-3, SWSN-2.2, SWSN-3, SWSN-6, and PHF-10), collectively referred to as common factors (46,54). These common factors are bound by assembly-specific subunits in a mutually exclusive manner, which confers the distinct character of each of the two assemblies (**Fig 2 A**). Due to the absence of thorough biochemical investigation into the SWI/SNF complex in *C. elegans*, previous publications have classified subunits as part of the SWI/SNF core, accessory, or BAF/PBAF assemblies based on homology and phenotypic analyses (35,43,52,55). The prevailing model for the two SWI/SNF assemblies in *C. elegans* is that either the SWSN-8 subunit associates with common factors to form the BAF assembly, or the SWSN-7, SWSN-9, and PBRM-1 subunits associate with common factors to form the PBAF assembly (43,55,56). Prior investigations into SWI/SNF have revealed a wide array of developmental contexts in which the BAF and PBAF assemblies have overlapping and distinct roles in the regulation of cell cycle control, differentiation, and differentiated behavior (35,52,55,57–61).

**Figure 2.**
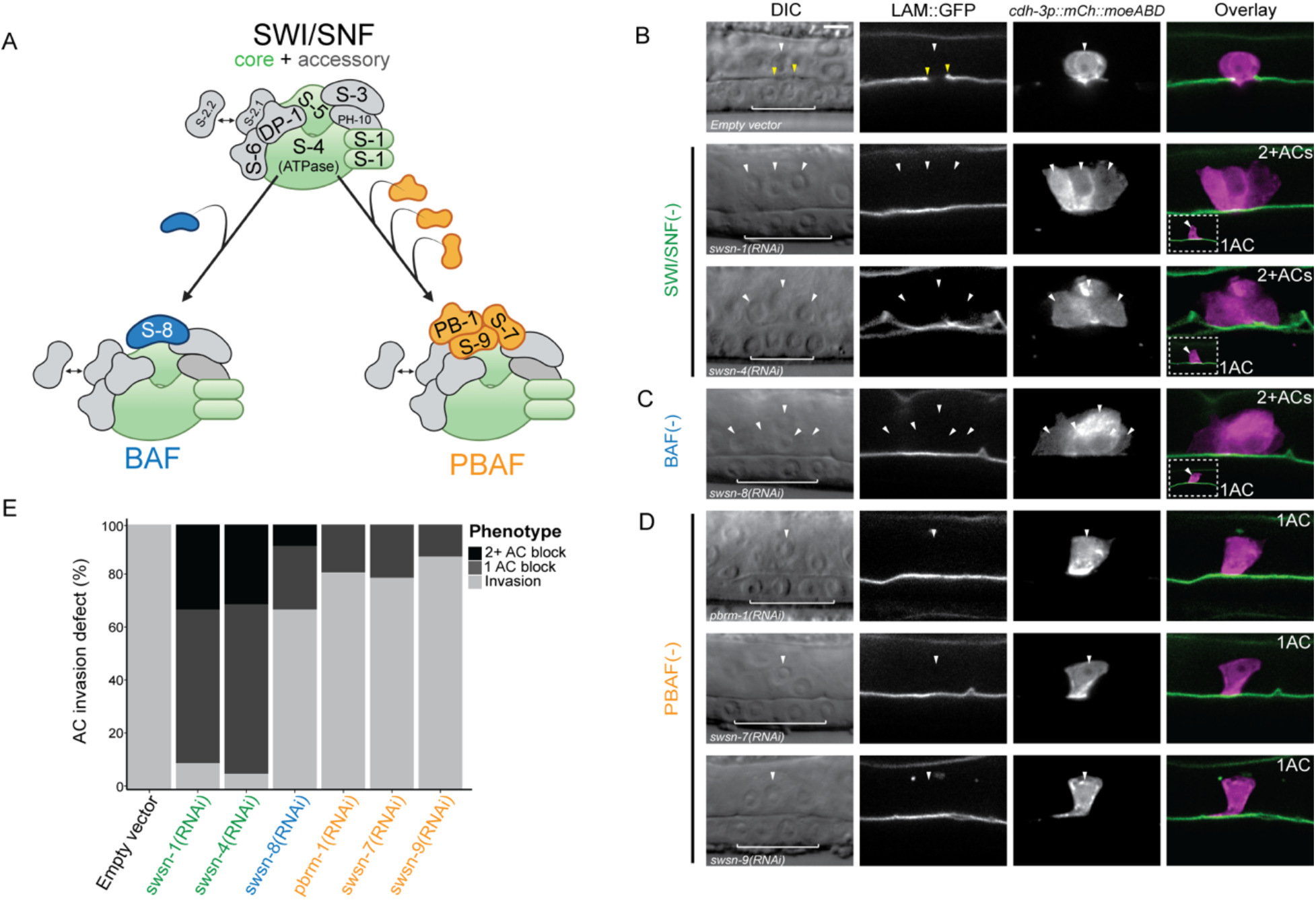
Enhanced RNAi targeting SWI/SNF core, BAF, and PBAF assembly subunits results in penetrant invasion defects. **(A)** Schematic depicting the *C. elegans* SWI/SNF core (middle, green) and accessory (middle, gray) subunits and BAF (left, blue), and PBAF (right, orange) assemblies. **(B-D)** DIC (left), corresponding fluorescence images (middle), and fluorescence overlay (right) representing loss of AC (magenta, *cdh-3p::mCherry::moeABD*) invasion through the BM (green, *laminin::GFP*) following RNAi depletion of SWI/SNF core (*swsn-1* and *swsn-4*) **(B)**, BAF (*swsn-8*) **(C)**, and PBAF (*pbrm-1, swsn-7,* and *swsn-9*) **(D)** subunits. In cases where multiple cells expressed the AC reporter (2+ACs) in the same animal, each is indicated with a white arrowhead. In cases where multiple cells expressed the AC reporter (2+ACs), a representative image from the same treatment of a single AC that fails to breach the BM is displayed as an inset (white dashed box, bottom left). White arrowheads indicate AC(s), yellow arrowheads indicate boundaries of breach in BM, and white brackets indicate 1° VPCs. **(E)** Stacked bar chart showing the penetrance of AC invasion defects following treatment with SWI/SNF RNAi depletion, binned by AC phenotype (n≥50 animals examined for each treatment).

To investigate the contribution of individual SWI/SNF subunits to AC invasion and to distinguish potentially distinct roles of the BAF and PBAF assemblies, we generated improved RNAi constructs utilizing the T444T vector (62) to target representative subunits of the core and both SWI/SNF assemblies (**Table S3**). Knockdown of SWI/SNF subunits in whole-body RNAi sensitive animals following treatment with T444T RNAi vectors resulted in penetrant loss of invasion. The majority of ACs failed to invade following treatment with RNAi targeting the core SWI/SNF ATPase subunit *swsn-4* or core subunit *swsn-1* (90% and 94%, respectively; n=50 animals; **Fig 2 B, E**). RNAi-mediated knockdown of the BAF assembly subunit *swsn-8* also resulted in significant loss of AC invasion (32%; n=50 animals; **Fig 2 C, E**). Knockdown of the PBAF assembly subunits with *pbrm-1(RNAi), swsn-7(RNAi)*, or *swsn-9(RNAi)* resulted in a less penetrant loss of AC invasion (18%, 20%, and 12%, respectively; n=50 animals; **Fig 2 D, E**). Importantly, at least one cell in the ventral uterus expressed the fluorescent AC reporter in all animals across all treatments, suggesting that loss of the SWI/SNF complex does not preclude the birth of an AC.

Interestingly, in addition to a single non-invasive AC phenotype, RNAi-mediated knockdown of *swsn-1*, *swsn-4* or *swsn-8* also resulted in a second phenotype characterized by multiple uterine cells expressing the AC reporter (*cdh-3p::mCherry::moeABD*) which failed to invade the BM (*laminin::GFP*) (32%, 30% and 8%, respectively; **Fig 2 B-C, E**). In all instances where more than one cell expressed the AC reporter, no breach in the underlying BM was detected at the P6.p 4-cell stage. In contrast, only the single non-invasive AC phenotype resulted from RNAi treatment targeting PBAF assembly subunits (**Fig 2 D-E**). These results suggest that the SWI/SNF assemblies BAF and PBAF may promote AC invasion through distinct mechanisms, perhaps via regulation of both a cell cycle-dependent and -independent mechanism, respectively.

### Characterization of endogenous GFP reporter alleles and the efficacy of improved SWI/SNF RNAi vectors

Next, to confirm expression of SWI/SNF subunits in the AC and to quantitatively assess the potency of our enhanced SWI/SNF RNAi vectors we utilized CRISPR/Cas9 genome engineering to generate GFP-tagged alleles of *swsn-4* and *swsn-8*, inserting a codon-optimized GFP tag into the 5’ end and 3’ end of the *swsn-4* and *swsn-8* loci, respectively (**Fig S2 A**) (63). The GFP-tagged endogenous strains showed ubiquitous and nuclear-localized expression of GFP::SWSN-4 and SWSN-8::GFP throughout the *C. elegans* developmental life cycle (**Fig S2 B**). We also obtained a strain containing an endogenously eGFP-labeled PBAF subunit (*pbrm-1::eGFP*) from the *Caenorhabditis* Genetics Center (CGC). We quantified fluorescence protein expression of SWI/SNF core ATPase (GFP::SWSN-4), BAF (SWSN-8::GFP), and PBAF (PBRM-1::eGFP) subunits in the AC during vulval development across the L3 and early L4 stages, as defined by the division pattern of the 1°-fated VPCs (14) (n≥28 animals per stage; **Fig S3 A-C’**). Expression of all three subunits was enhanced in the AC relative to the neighboring ventral uterine (VU; *swsn-4*: 18%, *swsn-8*: 21%, *pbrm-1*: 17% enhanced) and 1° VPC (*swsn-4*: 30%, *swsn-8*: 38%, *pbrm-1*: 23% enhanced) lineages during AC invasion (P6.p 2 cell – 4 cell stage, **Fig S3 A’-C’**). Late in vulval development at the P6.p 8 cell stage, expression of GFP::SWSN-4 and PBRM-1::eGFP increases in the 1° VPCs and is no longer statistically separable from expression in the AC, whereas expression of SWSN-8::GFP in the VPCs remains significantly lower than in the AC (**Fig S3 A’-C’**).

We treated SWI/SNF endogenously labeled GFP-tagged strains with our RNAi vectors to precisely quantify the efficiency of RNAi-mediated knockdown of target SWI/SNF complex subunits and to correlate this loss with the resulting AC phenotypes. Treatment with either *swsn-4(RNAi)* or *swsn-8(RNAi)* vectors resulted in robust depletion of fluorescence expression of GFP::SWSN-4 (94% depletion) and SWSN-8::GFP (81% depletion) in the AC (**Fig S4 A-B, D**) and penetrant loss of invasion (90% and 30%, respectively; n=30 animals for each condition; **Fig S4 E**). We also noted instances where multiple cells expressed the AC reporter (23% and 10%, respectively; n=30 animals for each condition; **Fig S4 E**). Treatment of the PBRM-1::eGFP strain with *pbrm-1(RNAi)* revealed weaker but significant knockdown of PBRM-1 protein (49% depletion), and a lower penetrance of invasion defects (17%; n=30 animals; **Fig S4 C-E**). It is unclear why PBRM-1::eGFP endogenous protein level in the AC of animals treated with enhanced *pbrm-1(RNAi)* remains considerably higher compared to treatment of strains containing *swsn-4* or *swsn-8* reporter alleles with their respective RNAis. We hypothesize that this may be the consequence of differential protein perdurance. Altogether, these results confirm the dynamic expression of the SW/SNF core, BAF, and PBAF subunits in the AC before, during, and after invasion and demonstrate the effectiveness of our improved SWI/SNF-targeting RNAi vectors.

### *C. elegans* SWI/SNF subunits exhibit low levels of intracomplex cross-regulation

Work in mammalian cell culture has revealed that the mSWI/SNF complex is assembled in a step-wise fashion, with stability of the complex as a whole and association of individual subunits depending on the prior expression and association of other subunits (64). To date it is unknown whether in *C. elegans* individual SWI/SNF subunits activate other SWI/SNF subunits. It is also unclear whether subunits of the two assemblies in *C. elegans* – BAF and PBAF – stabilize the core protein subunits or vice-versa. Therefore, we used our endogenously labeled GFP-SWI/SNF strains to ask whether transcriptional knockdown of individual subunits of the core, BAF, or PBAF induce changes in protein expression of other subunits at the time of AC invasion (**Fig S5**). First, to determine whether representative subunits of the SWI/SNF assemblies promote or stabilize the ATPase of the complex, we treated *GFP::swsn-4* animals with either *swsn-8(RNAi)* or *pbrm-1(RNAi)* (**Fig S5 A**). Quantification of fluorescence expression in AC nuclei of *swsn-8(RNAi)* treated animals at the P6.p 4 cell stage revealed significantly lower GFP::SWSN-4 levels relative to the control group (34% GFP::SWSN-4 depletion; **Fig S5 A, D**). RNAi knockdown of the PBAF subunit *pbrm-1* also resulted in a significant but weaker loss of ATPase expression in the AC (11% GFP::SWSN-4 depletion; **Fig S5 A, D**). These results suggest that individual subunits of either SWI/SNF assembly may contribute to the protein stability and/or expression of the SWI/SNF ATPase in the *C. elegans* AC, with the BAF complex playing a potentially dominant activating role with respect to the ATPase.

Next, we treated animals containing either the *swsn-8* or *pbrm-1* endogenous GFP-reporters with enhanced RNAi to knockdown the expression of the SWI/SNF ATPase or the representative subunit of the alternative SWI/SNF assembly. Interestingly, while unaffected by knockdown of the PBAF assembly subunit *pbrm-1,* RNAi knockdown of the ATPase *swsn-4* resulted in a 42% increase in the expression of SWSN-8::GFP in the AC (**Fig S5 B, D**). Finally, relative to the expression of the endogenous PBAF subunit in the ACs of animals treated with the empty vector control RNAi, AC nuclei of PBRM-1::eGFP animals treated with *swsn-4(RNAi)* had significantly lower levels of protein expression (38% PBRM-1::eGFP depletion), whereas ACs in *swsn-8(RNAi)* treated animals expressed 13% more PBRM-1::eGFP (**Fig S5 C-D**).

Since knockdown of either *swsn-4* or *swsn-8* subunits resulted in two distinct phenotypes – individual animals with single non-invasive ACs and animals with multiple non-invasive cells expressing the AC-reporter - we next sought to determine whether these two phenotypes were distinct with respect to SWI/SNF subunit expression. To do this, we binned data from the intracomplex RNAi experimental series (above) into the two non-invasive phenotypes and compared the fluorescence expression levels of the endogenous proteins within SWI/SNF RNAi conditions. Given the infrequency of the multi-AC phenotype, statistical comparisons were necessarily limited to treatments in which the population of animals contained at least 10 multi non-invasive ACs. Treatment of SWSN-8::GFP with *swsn-4(RNAi)* resulted in a total of 24 multi non-invasive ACs (53 ACs total; n=41 animals) and no significant difference was detected in SWSN-8::GFP expression between the nuclei of the single non-invasive AC phenotype and the multi non-invasive AC phenotype groups (**Fig S5 E**). The second statistical comparison was made between the two phenotypes in PBRM-1::eGFP animals treated with *swsn-8(RNAi)* (**Fig S5 E**), in which 14 multi non-invasive ACs were detected (51 ACs total; n=42 animals). Quantification of endogenous PBRM-1::eGFP fluorescence expression in this condition revealed a slight (12%) increase in expression of the PBAF subunit in the nuclei of ACs of the multi non-invasive phenotype group compared to the single non-invasive phenotype (**Fig S5 E**).

Based on these results, a tentative model for epistatic interactions between the SWI/SNF ATPase, BAF, and PBAF assembly subunits can be composed for the AC (**Fig S5 F**). Our data indicate that some degree of SWI/SNF intra- and inter-complex regulation occurs in the AC. We find that the most significant aspect of SWI/SNF intra-complex regulation is exercised by the ATPase on the assembly specific subunits, where *swsn-4* knockdown results in a significant increase in BAF/SWSN-8 and a significant decrease PBAF/PBRM-1. SWI/SNF intercomplex regulation appears to be weaker in the AC as knockdown of *pbrm-1* does not affect SWSN-8::GFP expression, and knockdown of *swsn-8* results in a slight increase in PBRM-1::GFP expression.

### The SWI/SNF ATPase SWSN-4 provides dose-dependent regulation of AC invasion

The degree to which the SWI/SNF complex contributes to tumorigenesis in clinical settings has been linked to the dose of functional SWI/SNF ATPase in precancerous and transformed cells (32,34,65). Previous work in *C. elegans* has demonstrated a similar dose dependent relationship between SWI/SNF and cell cycle control (35). To determine whether the phenotypic dosage sensitivity seen in cancer and *C. elegans* mesodermal (M) cell development is also characteristic of SWI/SNF in the promotion of AC invasion (30), we modulated expression of GFP::SWSN-4 using a combination of RNAi-mediated knockdown and AC-specific GFP-targeting nanobody technology.

Though RNAi treatment targeting the *swsn-4* subunit in the endogenously-tagged strain resulted in significant knockdown of fluorescence expression of GFP::SWSN-4 in the AC, loss of expression was noted in many other tissues in treated animals, including the 1° VPCs, which contribute to AC invasion non-autonomously (14,66) (**Fig S4**). Thus, to limit loss of expression to the AC, we used an anti-GFP nanobody fused to a SOCS-box containing a ubiquitin ligase adaptor, driven with tissue-specific promoters to achieve lineage-restricted protein depletion (30) (**Fig 3**). To follow the expression of the anti-GFP nanobody transgenes, we also included a fluorescent histone label separated from the anti-GFP nanobody sequence by the p2a viral self-cleaving peptide (*ACp::antiGFP-nanobody::p2a::His-58::mCherry*). We generated two anti-GFP nanobody constructs, using conserved *cis*-regulatory elements from the *cdh-3* and *egl-43* promoters (22,24,39,67,68) and introduced them into a strain containing the endogenous GFP-tagged allele of *swsn-4* as well as background AC and BM reporters (**Fig 3 B-C**). The *cdh-3*-driven nanobody transgene (*cdh-3p::antiGFP-nanobody::p2a::his-58::mCherry*) resulted in a weak reduction of GFP::SWSN-4 levels with no significant difference in fluorescence expression in the AC compared to wildtype animals (6% depletion; n=80 animals; **Fig 3 F**); however, consistent with the wildtype expression of the *cdh-3* promoter (22,39), it expressed specifically in the AC and resulted in defective AC invasion, suggesting partial loss of function (20.6% AC invasion defect; n=102; **Fig 3 B, G**). The *egl-43p::antiGFP-nanobody* transgene (*egl-43p::antiGFP-nanobody::p2a::His-58::mCherry*) expression pattern was also consistent with the wildtype expression characterized in previous work (22,67–69), as indicated by nuclear expression of HIS-58::mCherry in the AC and in the neighboring ventral uterine and dorsal uterine (VU/DU) cells (**Fig 3 C**; asterisk denotes HIS-58::mCherry expression in a non-AC ventral uterine cell) (22,39,67). Importantly, as the AC invades independent of VU/DU cells (14), anti-GFP expression in these tissues should not affect AC invasion. Similar to animals treated with *swsn-4(RNAi)* (**Fig 3 D**), *egl-43p::antiGFP-nanobody*-mediated protein depletion of GFP::SWSN-4 resulted in a significant loss of fluorescence expression in the AC (71% GFP depletion; n=80 animals; **Fig 3 C, F**) as well as a penetrant loss of invasion and incidence of individual animals with multiple uterine cells that were in contact with the ventral BM and expressed the AC reporter (88.2% AC invasion defect, 2.9% multiple AC phenotype; n=101 animals; **Fig 3 G**). These results support our uterine-specific SWI/SNF RNAi results and provide strong evidence for a cell-autonomous role for the SWI/SNF complex in promoting cell invasion and cell cycle arrest.

**Figure 3.**
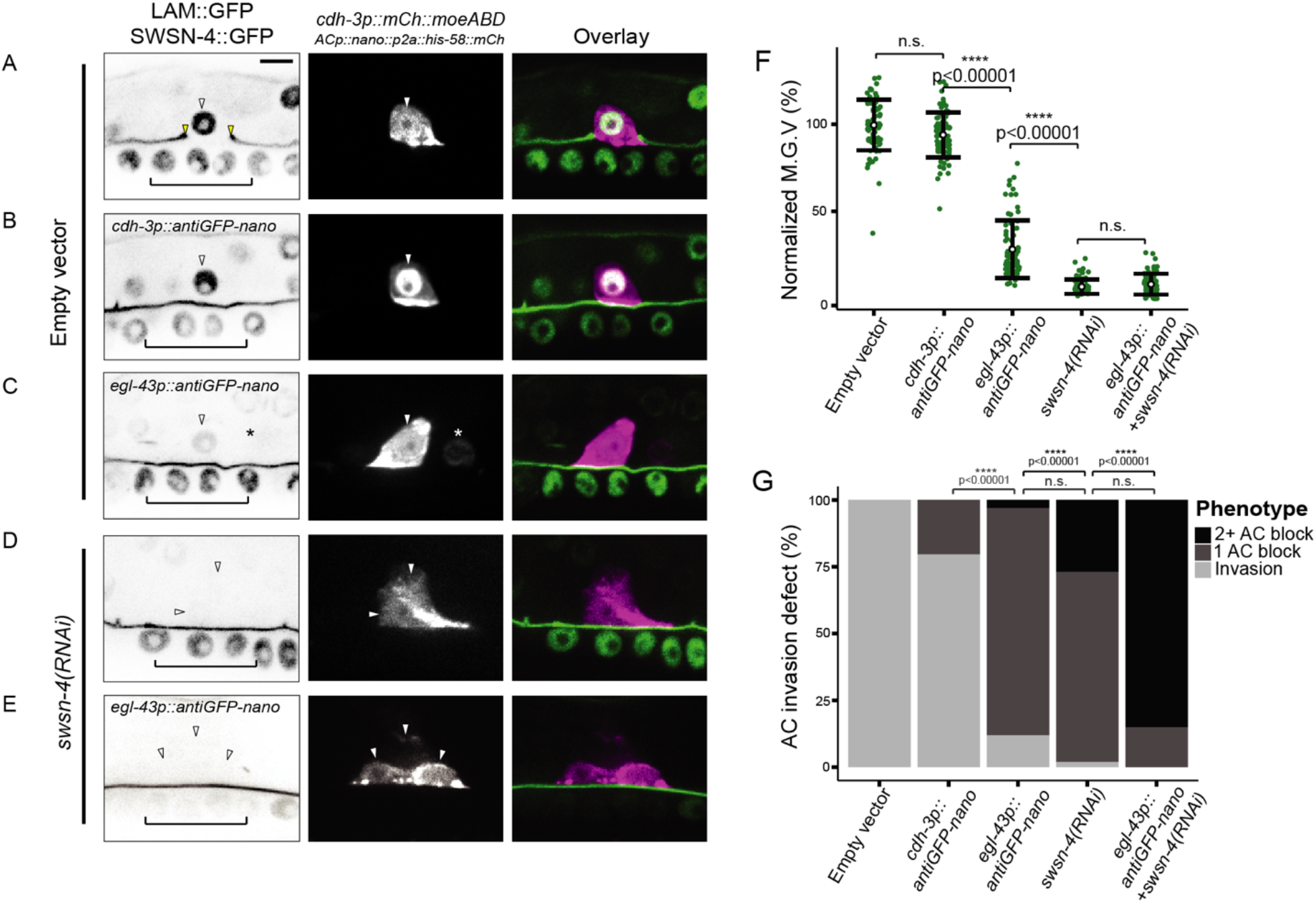
AC invasion and cell cycle arrest depend on dosage of SWI/SNF ATPase. **(A-E)** Representative fluorescence images depicting expression of BM marker (*laminin::GFP*) and endogenous GFP::SWSN-4 (left), AC reporter (*cdh-3p::mCherry::moeABD*, middle), and fluorescence overlay (right) across experimental treatments. White arrowheads indicate AC(s), yellow arrowheads indicate boundaries of breach in BM, and black brackets indicate 1° VPCs. In cases where multiple cells expressed the AC reporter in the same animal, each is indicated with a single white arrowhead. Asterisk indicates anti-GFP nanobody expression in neighboring VU cell. **(F)** Quantification of mean gray values (M.G.V.) of endogenous GFP::SWSN-4 in ACs in control animals (empty vector) and across all experimental treatments normalized to mean fluorescent expression in wildtype animals (n≥40 animals per treatment, p values for Student’s *t*-test comparing expression of successive knockdown are displayed on the figure). In this and all other figures, open circles and error bars denote mean±standard deviation (s.d.). n.s. not significant. **(G)** Stacked bar chart showing quantification of AC invasion defects corresponding to each treatment, binned by AC phenotype (n≥40 animals per condition; p values for Fisher’s exact test comparing phenotypes of successive knockdown strategies are displayed above compared groups). Grey brackets indicate statistical significance between invasion total in each condition compared to invasion defect total. Black brackets indicate statistical significance between incidences of invasion defects with multiple ACs compared to incidences of invasion defects with single ACs. n.s. not significant.

To further deplete *swsn-4* expression in the AC, we treated transgenic *egl-43p::antiGFP-nanobody* animals with *swsn-4(RNAi)* (**Fig 3 E**). Strikingly, in this combination knockdown strategy, 100% of AC invasion was lost and the frequency of multiple cells expressing the AC specification reporter drastically increased relative to treatment with *swsn-4(RNAi)* or the *egl-43*-driven anti-GFP nanobody conditions alone (83% multiple AC phenotype; n=41 animals; **Fig 3 G**). Together, these results demonstrate a phenotypic spectrum that corresponds to successive loss of *swsn-4* in the AC: whereas moderate loss of the ATPase results in single non-invasive ACs in animals containing *egl-43p::antiGFP-nanobody*, strong loss of expression in the *egl-43p::antiGFP-nanobody* background or following treatment with *swsn-4(RNAi)* results in animals with both single and multiple non-invasive ACs; in the strongest knockdown condition – *egl-43p::antiGFP-nanobody* animals treated with *swsn-4(RNAi)* - multiple non-invasive ACs were present per animal with near complete penetrance. Though the combination of *swsn-4(RNAi)* and antiGFP-nanobody-mediated depletion resulted in robust loss of expression of the core ATPase of the SWI/SNF complex, the fluorescence expression was not significantly different than treatment with *swsn-4(RNAi)* alone (93% vs. 92% GFP depletion, respectively; n≥41 animals for each treatment; **Fig 3 G**); therefore we theorize that the fluorescence values were beyond our threshold ability to quantify based on the fluorescence detection limits of our imaging system. Altogether, our data demonstrated that in the AC, the ATPase of the SWI/SNF complex contributes to invasion cell-autonomously and in a dose-dependent manner: moderate loss of expression produced non-invasive single ACs while extreme loss of expression led to a non-invasive hyperproliferative state.

### Improved *swsn-4(RNAi)* vector is sufficient to recapitulate null phenotype in M lineage

A recent study focusing on cell cycle control of SWI/SNF throughout *C. elegans* muscle and epithelial differentiation demonstrated tissue and lineage-specific phenotypes following weak or strong loss of core SWI/SNF subunits (32). Within the M lineage that gives rise to posterior body wall muscles (BWMs), coelomocytes (CCs), and reproductive muscles or sex myoblast (SMs) descendants, different cell types responded differently to loss of SWI/SNF. In the BWM, strong loss of SWI/SNF resulted in hyperproliferation, like the phenotype we detect in the AC. The opposite is true in the SM lineage, where modest knockdown of *swsn-4* resulted in hyperproliferation while complete loss of *swsn-4* expression resulted in a null phenotype where SMs failed to divide and arrest in S phase (35). We next sought to validate the strength of our enhanced *swsn-4(RNAi)* vector by examining proliferative state in the SMs. To accomplish this, we treated animals containing a lineage-restricted cyclin-dependent kinase (CDK) activity sensor (*unc-62p::DHB::2xmKate2*) with *swsn-4(RNAi)* (**Fig S6 A**). In this genetic background, we determined the number (**Fig S6 B**) and cell cycle state (**Fig S6 C**) of SM cells at a time when the majority of SMs in control animals had finished cycling and subsequently differentiated (late P6.p 8 cell stage; 16 SM cell stage). Animals treated with *swsn-4(RNAi)* had significantly fewer SM cells than controls (mean SMs/animals = 5; n=31 animals; **Fig S6 B**) with many instances of SMs that failed to enter a single round of cell division (n=20 single SMs out of 43 animals). Interestingly, 28% (12/43) of animals treated with *swsn-4(RNAi)* were absent of SMs on either the left or the right side, whereas 100% (30/30) control animals had SMs on both sides, which may indicate a defect in specification, early cell division, and/or migration of SMs. To quantify cell cycle state, we measured localization of an SM-specific CDK sensor, which uses a fragment of mammalian DNA Helicase B (DHB) fused to two copies of mKate2 (36,70). In cells with low CDK activity that are quiescent or post-mitotic, the ratiometric CDK sensor is strongly nuclear localized (36,68,70). In cycling cells with increasing CDK activity, the CDK sensor progressively translocates from the nucleus to the cytosoplasm, with a ratio approaching 1.0 in S phase and >1 in cells in G_2_ (36). Thus, the cytoplasmic:nuclear (C/N) ratio of DHB::2xmKate2 can serve as a proxy to identify cell cycle state. By the time the majority of SMs in the control condition were differentiating and arrested in a G_0_ cell cycle state (mean C/N ratio=0.320; n=90 SMs; **Fig S6 C**), many animals treated with *swsn-4(RNAi)* had single SMs that failed to divide and a mean DHB C/N ratio indicative of a long pause or arrest in S phase (36) (Avg. C/N ratio = 0.803; n=20 SMs; **Fig S6 C**). Together, these results suggested that the strength of our enhanced *swsn-4(RNAi)* targeting vector is sufficient to recapitulate the *swsn-4* null condition in the SM lineage, as we detected both the hypoproliferative phenotype and S-phase arrest that was observed using a lineage-restricted catalytically inactive SWI/SNF ATPase (35).

### The BAF assembly contributes to AC invasion via regulation of G_0_ cell cycle arrest

Having established that strong depletion of the SWI/SNF complex results in a fully penetrant defect in AC invasion with a high percentage of individual animals possessing multiple non-invasive ACs (**Fig 3**), we next investigated whether the extra ACs observed were the consequence of inappropriate AC proliferation (22,68). To determine whether the SWI/SNF complex is required for G_0_/G_1_ cell cycle arrest in the AC, we quantified CDK activity in the AC using a ubiquitously expressed *rps-27p::*DHB::GFP transgene paired with AC (*cdh-3p::mCherry::moeABD*) and BM (*laminin::GFP*) reporters in live animals following RNAi-mediated knockdown of SWI/SNF core (*swsn-4*), BAF (*swsn-8*), and PBAF (*pbrm-1*) subunits (**Fig 4**). In wild-type invasive ACs, we observed strong nuclear localization of the CDK sensor and quantified a cytoplasmic/nuclear (C/N) ratio indicative of G_0_/G_1_ arrest (mean C/N ratio: 0.226+/-0.075, n=67 animals) (**Fig 4 A, F**). In order to distinguish whether the wild-type AC C/N ratio is actually indicative of G_0_ rather than G_1_ arrest, we quantified the CDK activity in the neighboring uterine Pi cells at the P6.p 8 cell 1° VPC stage following their terminal division to establish a G_0_ reference point (71,72) (mean C/N ratio: 0.206+/-0.078, n=30 animals) (**Fig 4 E-F**). We found no significant difference between the CDK activity of terminal Pi cells and wild-type invading ACs, suggesting that the wild-type AC exists in a CDK^low^ G_0_, pro-invasive state (**Fig 4 A, E-F**). In animals treated with *pbrm-1(RNAi)*, the CDK sensor also localized principally in the nucleus of ACs that failed to invade (mean C/N ratio: 0.157+/-0.063, n=41 animals) and only a single non-invasive AC was observed per animal (**Fig 4 D, F**). In contrast, following treatment with *swsn-8*(*RNAi*) the majority of ACs that failed to invade the BM were in the G_1_/S phases of the cell cycle (mean C/N ratio: 0.636+/-0.204, n=21 animals) (**Fig 4 C, F**). Finally, like the *swsn-8(RNAi)* condition, loss of expression of the core ATPase of the SWI/SNF complex through treatment with *swsn-4*(*RNAi*) resulted in a broad range of C/N ratios (C/N ratio min: 0.240, C/N ratio max: 1.140, mean C/N ratio: 0.566+/-0.205; n=40 animals) in animals with single or multiple non-invasive ACs (**Fig 4 B, F**). Interestingly, the *swsn-4(RNAi)* treatment resulted in a higher proportion of non-invasive G_0_ phase (C/N ratio < 0.3) ACs (14%, n=48 cells) than were present in the *swsn-8(RNAi)* treated population (8%, n=25) (**Fig 4 B, E**). This finding reemphasizes the dependence of both SWI/SNF BAF and PBAF assemblies on the core ATPase of the complex, as the distribution of cell cycle states in ACs following *swsn-4(RNAi)* treatment represents both the cell cycle-dependent and cell cycle-independent phenotypes seen in ACs deficient in the *swsn-8* and *pbrm-1* subunits, respectively

**Figure 4.**
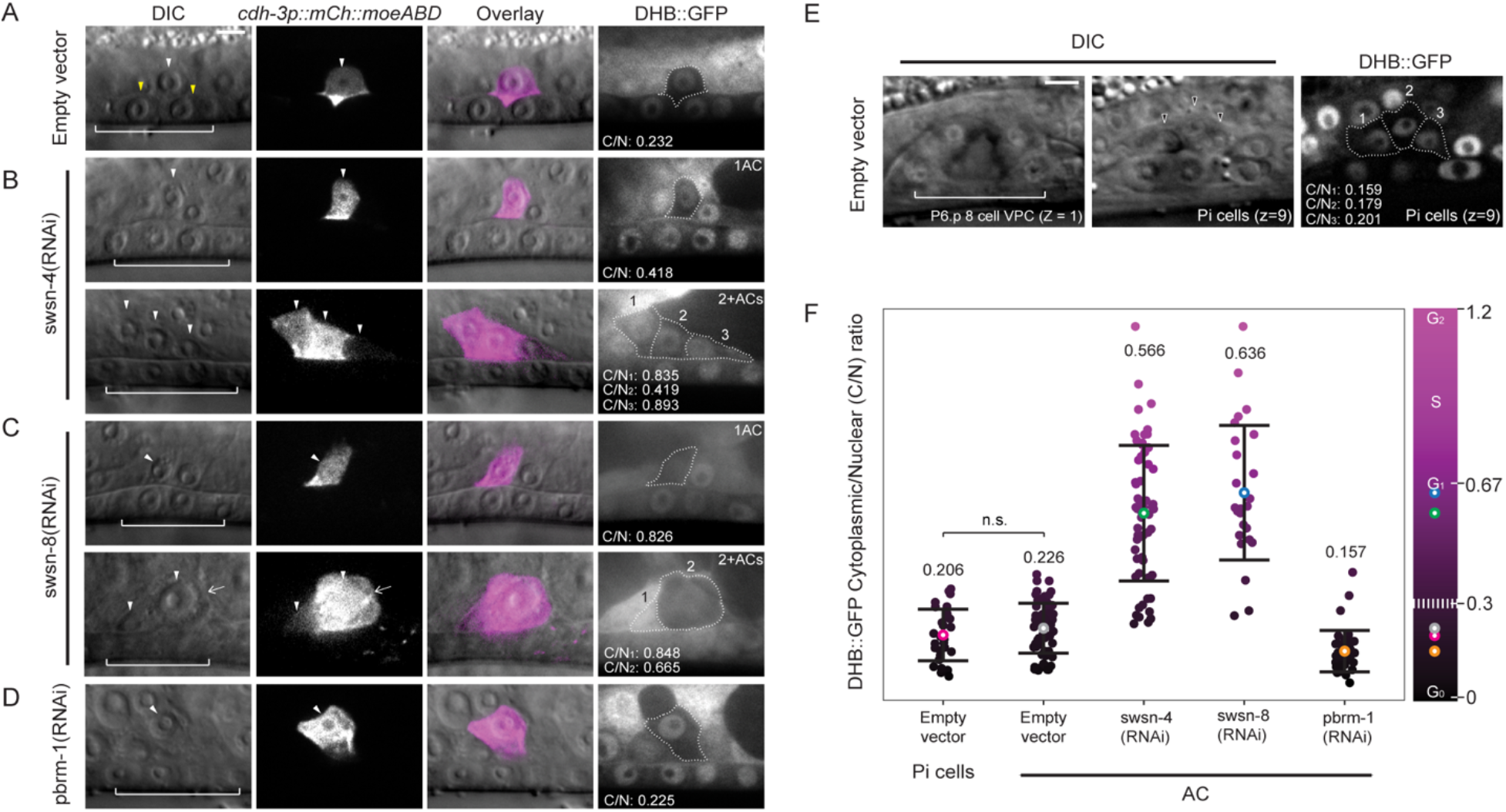
CDK sensor reveals SWI/SNF contribution to G_0_ arrest in the AC. Micrographs depicting DIC (left), AC (*cdh-3p::mCherry::moeABD*, center-left), DIC overlay (center-right), and DHB-based CDK activity sensor (right) in empty vector **(A)** and following treatment with SWI/SNF RNAi targeting subunits of the core (*swsn-4***, B)**, BAF (*swsn-8***, C)** and PBAF (*pbrm-1*, **D)** assemblies. White arrowheads indicate AC(s), yellow arrowheads indicate boundaries of breach in BM, and white brackets indicate 1° VPCs. In cases where treatment resulted in multiple cells expressing the AC reporter in the same animal, representative images of both single (1AC) and mitotic (2+ACs) phenotypes are given, and each AC is indicated with a single white arrowhead. Quantification of the cytoplasmic:nuclear (C/N) ratio of DHB::GFP in ACs (white dotted outline) is listed in the bottom left of each panel. Mitotic ACs are numbered, and C/N ratios are provided for each **(B-C)**. White arrowheads indicate AC(s), yellow arrowheads indicate boundaries of breach in BM, and white brackets indicate 1° VPCs. White arrow in C indicates an AC that is out of the focal plane. **(E)** Representative single z-plane micrographs of the vulva at the P6.p 8 cell stage (left, z=1) and the terminal Pi cells (middle, z=9) in DIC, and DHB-based CDK activity sensor in Pi cells (right). Quantification of the C/N ratio of DHB::GFP in three of four Pi cells (white dotted outline) that are in the plane of the image is listed in the bottom left. **(F)** Quantification of C/N DHB::GFP ratios for wild-type terminally divided Pi cells and all ACs in empty vector control and each RNAi treatment (n≥30 animals per treatment). Statistical comparison was made between the mean C/N ratio of ACs in control (empty vector) compared to control (empty vector) Pi cells using Student’s *t*-test (n≥30 for each stage and subunit; p values are displayed above compared groups). Mean C/N ratio is represented by colored open circles and correspond to numbers above the data. Gradient scale depicts cell cycle state as determined by quantification of each Pi cell or AC in all treatments (n≥30 animals per treatment), with dark/black depicting G_0_ and lighter/magenta depicting G_2_ cell cycle states. Dashed white line on gradient scale bar (right) corresponds to boundaries between G_0_ and G_1_ phases. Colored open circles on the gradient scale correspond to the mean C/N ratio in each of the same color. n.s. not significant.

### Forced G_0_ arrest through ectopic CKI-1 rescues invasive potential in BAF-deficient but not PBAF-deficient ACs

We have previously proposed and characterized a dichotomy that exists between invasion and proliferation in the AC (22,24). As evidence of this, loss of two of the three TFs that function in a cell cycle-dependent manner to maintain the AC in a cell cycle-arrested state (*nhr-67*/Tlx and *hlh-2*/Daughterless) can be rescued through induced expression of a cyclin dependent kinase inhibitor, *cki-1* (p21/p27) (22). These results suggest that, at least in some cases, TF activity can be bypassed completely to promote AC invasion by maintaining G_0_ arrest through direct cell cycle manipulation. To determine the extent to which the BAF assembly contributes to AC invasion through regulation of cell cycle arrest, we used a heat-shock inducible transgene to ectopically express CKI-1::mTagBFP2 in SWI/SNF-deficient ACs (**Fig 5**). Since the heat shock inducible transgene is ubiquitous and expresses variably between different animals and different tissues within an individual animal, we limited our analysis to animals with ACs with obvious mTagBFP2 fluorescence expression. While forced arrest in G_0_ was insufficient to significantly rescue AC invasion in animals treated with *swsn-4(RNAi)* (**Fig 5 B, B’, E**) or *pbrm-1(RNAi)* (**Fig 5 D, D’, E**), ectopic *cki-1* (CKI-1::mTagBFP2) expression in the AC significantly rescued cellular invasion in animals treated with *swsn-8(RNAi)* (**Fig 5 C, C’, E**). Strikingly, in 85.7% (6/7) of cases where ACs had proliferated prior to ectopic CKI-1 expression, forced G_0_ arrest led to multiple ACs breaching the BM (**Fig 5 F**), a phenotype we have reported previously using CKI-1 overexpression paired with loss of NHR-67. This demonstrated that mitotic ACs maintain the capacity to invade if they are re-arrested into a G_0_ state (24). To corroborate our CKI-1 heat shock data, we used an AC-specific CKI-1 transgene (*cdh-3p::CKI-1::GFP*) to induce G_0_ cell cycle arrest in *swsn-4-* and *swsn-8-*depleted ACs (**Fig S6**). Similar to the heat shock results, lineage-restricted expression of CKI-1::GFP failed to rescue invasion in animals deficient in *swsn-4* (**Fig S7 A-B**). However, transgenic *cdh-3p::CKI-1::GFP* animals treated with *swsn-8(RNAi),* had invasion defects significantly lower than control animals treated with *swsn-8(RNAi)* which lacked the G_0_ rescue transgene (**Fig S7 B**). Altogether, these data corroborate our DHB-based CDK sensor data (**Fig 4**), suggesting that the SWI/SNF assemblies differentially contribute to AC invasion with BAF specifically required for G_0_ cell cycle arrest.

**Figure 5.**
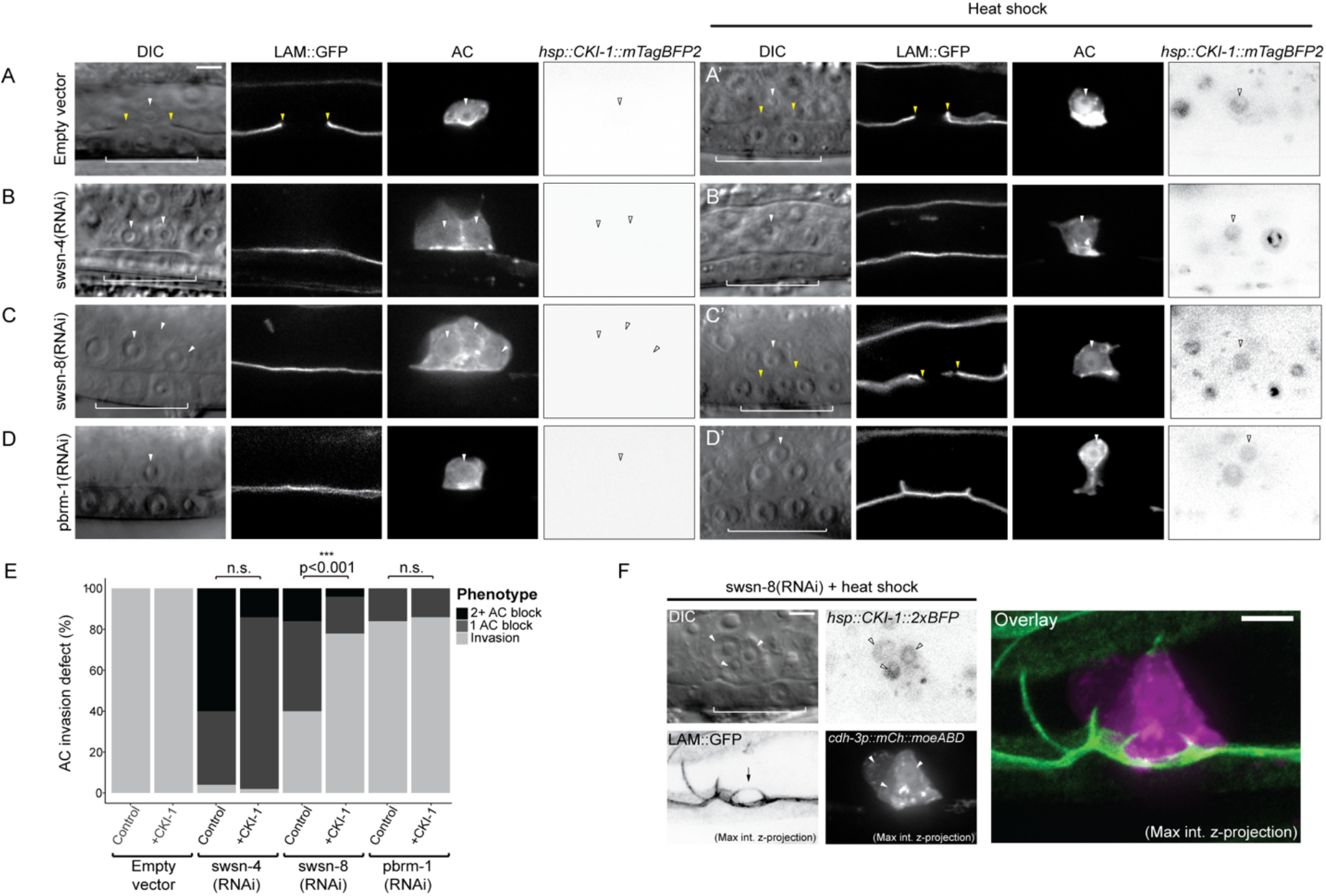
BAF depletion is rescued by G_0_ arrest. Representative micrographs depicting DIC (left), BM (*laminin::GFP*, center-left), AC (*cdh-3p::mCherry::moeABD,* center-right), and CKI-1 (*hsp::CKI-1::mTagBFP2*) expression in empty vector control **(A-A’)** and treatment with SWI/SNF RNAi under standard conditions **(A-D)** and following heat shock induction of CKI-1 **(A’-D’)**. CKI-1 images have been inverted for ease of visualization. White arrowheads indicate AC(s), yellow arrowheads indicate boundaries of breach in BM, and white brackets indicate 1° VPCs. **(E)** Stacked bar chart showing percentage of AC invasion defects corresponding to each RNAi treatment under standard growth conditions (control) and following heat shock induction of CKI-1 (+CKI-1), binned by AC phenotype (n≥30 animals per condition; Fisher’s exact test compared CKI-1(+) animals with control, non-heat shocked animals; p value is displayed above compared groups). n.s. not significant. **(F)** Representative micrographs of invasive group of *swsn-8* deficient ACs following induction of G_0_/G_1_ arrest. DIC (top-left), BM (bottom-left), CKI-1 expression (top-right), AC reporter (bottom-right). Max intensity z-projection of AC and BM reporter channels (bottom). Large breach in BM is indicated by black arrow.

### SWI/SNF chromatin remodeling promotes the invasive GRN in the AC

Previous work has demonstrated that the gene regulatory network (GRN) that promotes AC invasion consists of both cell cycle-dependent and cell cycle-independent TF subcircuits (22,68) (**Fig 1 B**). In the cell cycle-dependent subcircuit of the TF-GRN, *egl-43* (EVI1/MEL), *hlh-2* (E/Daughterless), and *nhr-67* (TLX/Tailless) cooperate in a type 1 coherent feed-forward loop that is reinforced via positive feedback to retain the AC in a post-mitotic, invasive state (22,68). The cell cycle-independent subcircuit of the AC TF-GRN is governed by the *fos-1* (FOS) TF with feedback from both *egl-43* and *hlh-2* (22). Since transcriptional knockdown of SWI/SNF ATPase results in both single and mitotic non-invasive AC phenotypes, we treated endogenously GFP-labeled strains for each TF in the GRN with *swsn-4(RNAi)* to determine whether SWI/SNF chromatin remodeling contributes to the regulation of either or both AC GRN subcircuits (**Fig 6**). In the cell cycle-dependent subcircuit, knockdown of the SWI/SNF ATPase resulted in significant loss of protein expression of EGL-43::GFP and NHR-67::GFP in the AC (39% and 26% GFP depletion, respectively; n≥41 animals; **Fig 6 A,C,E**). No significant difference was detected in the mean fluorescence expression of GFP::HLH-2 fusion protein in the AC upon knockdown of *swsn-4,* however the range of expression was broad following *swsn-4(RNAi)* treatment (∼2% GFP increase; n≥50 animals; **Fig 6 B, E**). In the cell cycle-independent subcircuit, loss of the SWI/SNF complex following treatment of *fos-1::GFP* animals with *swsn-4(RNAi)* resulted in a more moderate depletion of expression in the AC (11% GFP depletion; n≥50 animals; **Fig 6 D**). These results suggest that the SWI/SNF complex broadly remodels chromatin to promote both subcircuits of the pro-invasive AC GRN, though we are unable to determine whether the complex does so directly in regulatory regions of the lineage-specific pro-invasive TFs or in regulatory regions of genes that contribute to the regulation of the AC GRN.

**Figure 6.**
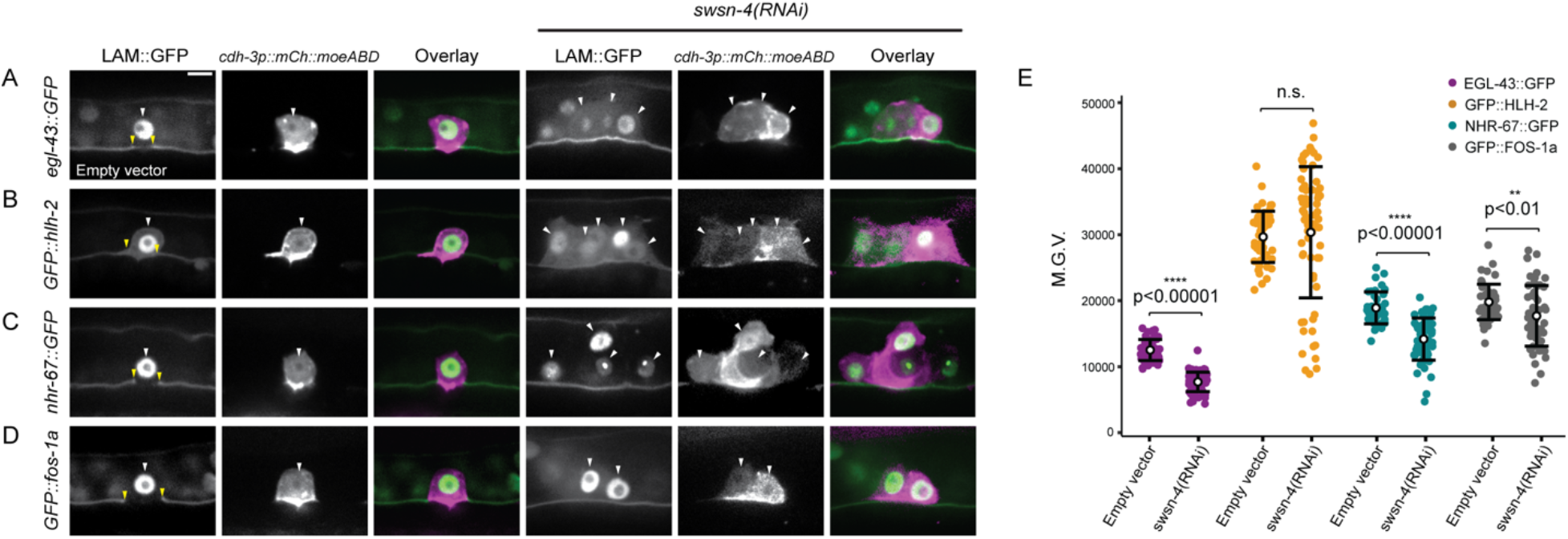
SWI/SNF regulates endogenously tagged TFs in the AC invasion GRN. Fluorescent micrographs depicting BM (*lam::GFP*) and AC (*cdh-3p::mCherry::moeABD*) expression of endogenously tagged TFs of the cell cycle-dependent subcircuit (*egl-43::GFP::egl-43* **(A),** *GFP::hlh-2* **(B),** and *nhr-67::GFP* **(C)**) and cell cycle-independent subcircuit (*GFP::fos-1a* **(D)**) of the AC GRN in animals treated with empty vector control (left) or *swsn-4(RNAi)* (right). White arrowheads indicate AC(s), yellow arrowheads indicate boundaries of breach in BM. **(E)** Quantification of fluorescent expression of each TF::GFP in ACs of control animals and animals treated with *swsn-4(RNAi)*. Statistical comparisons were made between the expression of each TF subunit in the AC in control and RNAi-treated animals using Student’s *t*-test (n≥30 for each condition; p values are displayed above black brackets). n.s. not significant.

### The PBAF assembly regulates AC contact with underlying BM

Previous investigations into SWI/SNF have demonstrated divergent roles for the PBAF assembly in cell cycle regulation. In yeast, Remodeling the Structure of Chromatin (RSC), the homologous complex to PBAF, is required for progression through mitosis (73,74). In *Drosophila,* the homologous complex PBAP does not appear to be required for mitotic progression; rather, cycling and G_2_/M transition is solely regulated by the BAF/BAP assembly (51). In the *C. elegans* M lineage, RNAi-mediated loss of BAF subunits results in hyperproliferation of the developing tissue, whereas knockdown of PBAF subunits has little effect on cell cycle control (35). Similarly, in this study, RNAi-mediated loss of PBAF subunits *pbrm-1*, *swsn-7*, or *swsn-9* resulted exclusively in single non-invasive cells expressing the AC reporter. However, given that the enhanced *pbrm-1(RNAi)* resulted in much weaker endogenous protein knockdown than the enhanced RNAis targeting either the SWI/SNF ATPase (*swsn-4*) or BAF assembly subunit (*swsn-8*) in the AC (**Fig S4**), and the dose-dependent phenotype following loss of the core ATPase (**Fig 3**), it is possible that we failed to observe the mitotic non-invasive AC phenotype due to insufficient PBAF subunit knockdown. To address this, we next asked whether *strong* loss of PBAF subunit expression contributes to the mitotic non-invasive AC phenotype. To accomplish this, we used an auxin inducible degron (AID)-RNAi combination knockdown strategy (75,76). We generated a strain with *pbrm-1* endogenously labeled with mNeonGreen and an auxin inducible degron (AID) (*pbrm-1::mNG::AID*) in a genetic background containing AC (*cdh-3p::mCherry::moeABD*) and BM (*laminin::GFP*) reporters. We then quantified fluorescence expression in the AC in this strain. When grown under standard conditions, 6.45% of the ACs had not invaded the BM by the P6.p 4 cell stage, suggesting a partial loss of function of *pbrm-1* (n=30; **Fig 7 A, E-F**). This partial loss of function phenotype is likely due to the insertion of the mNG::AID tag into the genomic locus, causing a putative hypomorphic allele. Next, we introduced a ubiquitous, mRuby-labeled TIR1 transgene (*eft-3p::TIR1::mRuby*) into the animals and assessed AC invasion under standard conditions (aux(-)) or in the presence of the auxin hormone (aux(+); **Fig 7 B**). We observed no statistically significant difference in the fluorescence expression of PBRM-1::mNG::AID protein in the AC, nor did we observe any differences in AC invasion defects between the strains with and without the TIR1 transgene when grown on aux(-) media (TIR1+: 3% depletion, 16.67% invasion defect; n=30; **Fig 7 E-F**). However, in both conditions, some ACs that invaded seemed to do so only partially, as most of the ventral epidermal BM remained intact beneath ACs (**Fig 7 B**, black arrowhead). In the aux(+) condition, there was a significant reduction in PBRM-1::mNG::AID protein level in the AC of animals containing the TIR1 transgene relative to the same strain grown in the aux(-) condition or the strain without the TIR1 transgene (49% and 51% depletion, respectively; n=30; **Fig 7 E**); however, there was no significant difference in the penetrance of AC invasion defects (16.67% invasion defect; n=30; **Fig 7 F**). Like our previous results with *pbrm-1(RNAi)* treated animals, we observed no extra cells expressing the AC reporter following loss of expression of PBAF in the AC using the AID system.

**Figure 7.**
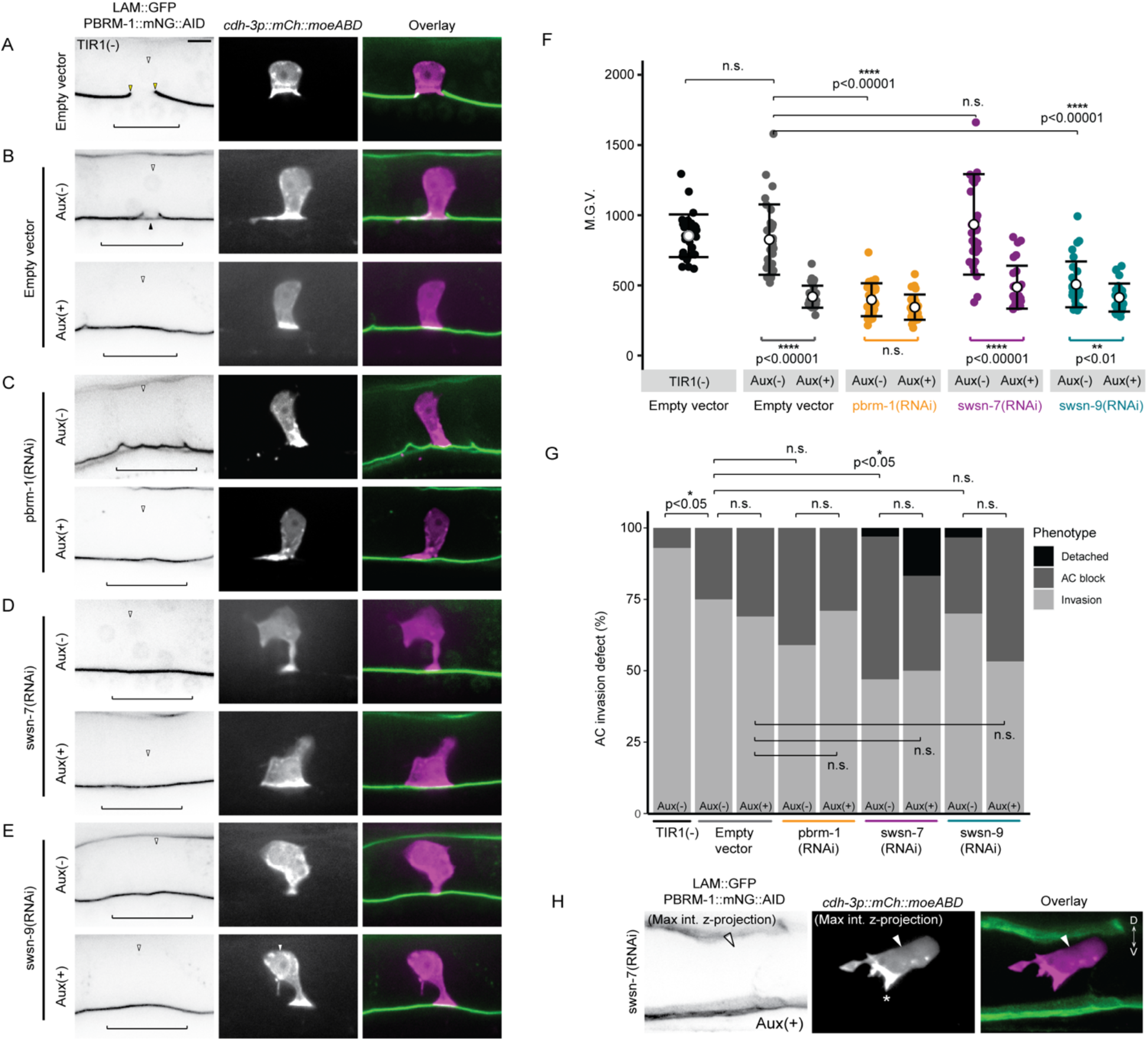
PBAF promotes AC contact to the BM. Representative micrographs of BM (*lam::GFP*) and endogenous *pbrm-1::mNG::AID* (left), AC (*cdh-3p::mCherry::moeABD*, center), and fluorescent overlays (right) of animals lacking **(A)** or possessing **(B-E)** ubiquitous TIR1 expression treated with empty vector control **(B)** or RNAi targeting PBAF subunits in the absence Aux(-) (top) or presence Aux(+) (bottom) of media containing the auxin hormone **(C-E).** PBRM-1::mNG::AID images have been inverted for ease of visualization. **(F)** Quantification of fluorescence expression (M.G.V) of PBRM-1::mNG::AID in ACs of animals in each condition (N≥30 animals in each treatment; p values for Fisher’s exact test comparing strains containing TIR1 to the TIR1(-) strain, and comparing strains containing TIR1 in the Aux(-) to the Aux(+) condition, are displayed above compared groups). **(G)** Stacked bar chart showing percentage of AC invasion defects corresponding to each treatment, binned by AC phenotype (N≥30 animals per condition; Fisher’s exact test compared penetrance of AC invasion defects between indicated conditions). Black brackets indicate statistical significance between invasion total in each condition compared to invasion defect total. **(H)** Max intensity z-projection of AC and BM reporter channels depicting a detached AC phenotype in *swsn-7*-deficient ACs in the Aux(+) condition. BM (left), AC (center), fluorescence overlay (right). Asterisk indicates polarized F-actin driven protrusion extending ventrally.

Next, we treated *pbrm-1::mNG::AID* animals containing ubiquitous TIR1 with *pbrm-1(RNAi)* in both aux(-) and aux(+) conditions. As expected, treatment of the *pbrm-1::mNG::AID* strain with *pbrm-1(RNAi)* resulted in very low expression of the subunit in the AC even in the absence of auxin and there was no significant difference in expression between the Aux(-) and Aux(+) conditions (**Fig 7 C,F**). Interestingly, there was also no significant difference in the penetrance of AC invasion defects between the *pbrm-1::mNG::AID* strain treated with control compared to the strain treated with *pbrm-1(RNAi)* in the presence of auxin (**Fig 7 G**). Since the combination treatment of a hypomorphic *pbrm-1* allele, Auxin-AID-mediated depletion of endogenous PBRM-1::mNG::AID, and *pbrm-1(RNAi)* does not result in a significant increase in AC invasion defects or non-invasive mitotic ACs, these results suggest that, unlike the dose-dependent contribution to invasion of *swsn-4*, the *pbrm-1* strong knockdown or null phenotype may be only partial/incomplete loss of AC invasion.

Since the PBAF assembly in *C. elegans* consists of several subunits, *pbrm-1* (PBRM1), *swsn-7* (ARID2), and *swsn-9* (BRD7/BRD9), we next investigated whether combinatorial knockdown of PBAF subunits would enhance the penetrance of AC invasion defects or result in the mitotic non-invasive AC phenotype. In the absence of auxin, there was no significant difference in PBRM-1::mNG::AID expression in the AC of animals treated with *swsn-7(RNAi)* compared to animals treated with empty vector control (n=30; **Fig 7 E**), however there was a significant increase in loss of AC invasion (50% invasion defect; n=30; **Fig 7 F**). Strikingly, in one case, the AC was completely detached from the BM, as we detected no AC membrane protrusions (*cdh-3p::mCherry::moeABD*) in contact with the ventral surface of the gonad (**Fig 7 F**). Animals treated with *swsn-7(RNAi)* and aux(+) had significantly lower expression of PBRM-1::mNG::AID in the AC when compared to animals treated with *swsn-7(RNAi)* in the aux(-) condition (49% depletion; n=30; **Fig 7 E**). While no significant difference was seen in loss of AC invasion in aux(+) (48.39% AC invasion defect), 16% (5/31) of animals in this treatment had ACs entirely detached from the ventral BM (n=31; **Fig 7 F-G**). In contrast to treatment with *swsn-7(RNAi)*, in the *swsn-9(RNAi)* aux(-) condition, PBRM-1::mNG::AID expression in ACs was significantly lower than that in the ACs of animals treated with empty vector control aux(-) (39% depletion; n=30; **Fig 7 E**). It is unclear why transcriptional knockdown of *swsn-9* specifically results in a decrease in PBRM-1 protein expression in the AC and we theorize this may be the result of a potential stabilizing interaction between the SWSN-9 and PBRM-1 proteins. We did detect a significant decrease in expression of PBRM-1::mN G::AID in ACs in *swsn-9(RNAi)* aux(+) compared to the *swsn-9(RNAi)* aux(-) condition (19% depletion; n=30; **Fig 7 E**), however, we saw no statistically significant difference in penetrance of AC invasion defects between the two conditions (30% vs. 43%; n=30; **Fig 7 F**). We also noted one animal with a detached AC in the *swsn-9(RNAi)* aux (-) condition and zero in the aux(+) condition (**Fig 7 F**). Importantly, we only observed one AC per animal across all combinatorial treatments, supporting the hypothesis that the PBAF assembly does not contribute to G_0_ cell cycle arrest in the AC.

Detached ACs in both the *swsn-7(RNAi)* and *swsn-9(RNAi)* AID combination knockdown conditions suggest that the PBAF assembly regulates AC contact with the ventral epidermal BM. A previous study has shown that AC attachment is regulated by the *fos-1/egl-43* cell cycle-independent subcircuit of the AC GRN via regulation of lamellipodin/*mig-10b* and non-autonomously via netrin/*unc-6* signaling (77). ACs deficient in components of this pathway are attached to the ventral epidermal BM when specified and gradually lose contact over time, with peak loss of contact occurring at the time of AC invasion at the P6.p 4-cell stage (77). In order to determine whether the PBAF assembly remodels chromatin to promote activation of this subcircuit of the AC GRN, we treated endogenously tagged *fos-1::GFP* (22) animals with *pbrm-1(RNAi)* and quantified fluorescence expression in ACs that displayed invasion defects (**Fig S8 A-B**). Animals treated with *pbrm-1(RNAi)* had a modest but statistically significant loss of FOS-1::GFP protein levels in non-invasive ACs (34% depletion; n=20; **Fig S8 B**), suggesting that the PBAF assembly partially regulates the *fos*-dependent pathway that mediates attachment to the underlying BM.

Since depletion of the PBAF assembly resulted in moderate loss of FOS-1::GFP in the AC, we next examined functional interactions between FOS-1 and PBRM-1. Given that the PBRM-1::mNG::AID allele was slightly hypomorphic, with ∼17% invasion defects in backgrounds with TIR1, we used the strain containing TIR1 as a sensitized background. We found that even without the addition of auxin, co-depletion with *fos-1(RNAi)* resulted in almost complete loss of AC invasion (96.77% invasion defect; n=31; **Fig S8 C-D**). Finally, we examined whether RNAi-mediated depletion of *pbrm-1* is synergistic with loss of downstream targets of FOS-1, the matrix metalloproteinases (MMPs). Previously, it has been shown that animals harboring null mutations for five of the six MMPs encoded in the *C. elegans* genome (*zmp-1,-3,-4,-5* and *-6*), show delayed AC invasion (21). RNAi depletion of *pbrm-1* in quintuple MMP mutants significantly and synergistically enhanced late invasion defects (scored at the P6.p 8-cell stage) in this background (24.24% invasion defect; n=33; **Fig S8 E-F**) as compared to loss of either *pbrm-1* (3.8%; n=52) or MMPs (0%; n=35) alone. Together, these results suggest that the PBAF assembly functions synergistically with FOS-1 to regulate AC invasion.

## DISCUSSION

### A tissue-specific CRF RNAi screen identified genes critical for cellular invasion

Previous work in the *C. elegans* AC and in cancer cell invasion has emphasized the necessity for dynamic chromatin states and chromatin regulating factors in the promotion of cellular invasion (24,79–83). In this study, we used the *Caenorhabditis elegans* AC invasion model as a single cell, *in vivo* system to identify a suite of CRFs that contribute to the process of cellular invasion. We performed a tissue-specific RNAi feeding screen to assess 269 genes implicated in chromatin binding, chromatin remodeling complexes, or histone modification. We do not claim that genes that we failed to identify as regulators of cellular invasion in the screen are unimportant for the process; however, RNAi-mediated loss of the majority of CRF genes in the screen did result in some penetrance of AC invasion defects (**Table S1**). This finding was expected, as many of the genes we screened are global regulators of the genome and broadly contribute to various cell biological processes. We extracted a list of the most penetrant regulators of cell invasion from the broader list (**Table S2**). Many genes and gene classes that we recovered as significant regulators of AC invasion are homologous to human genes that have been previously studied in the context of cellular invasion and tumorigenesis including *cec-6*/CBX1/CBX8 (82,84), *cfi-1*/ARID3A/ARID3C (85), *psr-1*/JMJD6 (86), *skp-1*/SNW1 (87), and several TAFs (*taf-*1/TAF1/TAF1L, taf*-5*/TAF5/TAF5L, *taf-7.1*/TAF7/TAF7L) (88–90). Additionally, we recovered nematode-specific genes including *nra-3, and cec-2,* and genes whose human homologs have not been previously studied in the context of cellular invasion to our knowledge, such as *cec-3* (homologous human protein is uncharacterized) and *gna-2*/GNPNAT. Since the majority of the CRF genes we identified as significant regulators of AC invasion have been previously studied in the context of invasion in human development and cancer metastasis, these results demonstrate the utility of the *C. elegans* AC invasion system as a genetically and optically tractable *in vivo* environment to corroborate and characterize previously identified CRFs that promote cellular invasion in human diseases such as rheumatoid arthritis and cancer. Future studies should continue to characterize the relationship between CRFs identified here and cellular invasion.

### The dose of SWI/SNF ATPase dictates the penetrance of defects in AC invasion

For the majority of this study, we focused on characterizing the contribution of the SWI/SNF ATP-dependent chromatin remodeling complex to cellular invasion as it was highly represented among our list of significant regulators of AC invasion (**Table S2**) and has been extensively studied in the context of both cellular invasion and cell cycle control across a variety of animal models and in human cancers (31,35,40,42,48,56,59,83,91–96). Prior whole-exome studies have determined that over 20% of human tumors harbor mutations in one or more subunits of the SWI/SNF complex (32,48,98). Among the most frequently mutated subunits of the chromatin remodeling complex throughout SWI/SNF-deficient cancers is the core ATPase subunit BRG1/SMARCA4 (48,99) and the mutually exclusive ATPase paralog to BRG (32,81,98,100).

Due to the high degree of structural similarity between BRG and BRM, mutation or epigenetic silencing of either ATPase can be compensated by expression of the other. Previous investigation has determined BRM to be an effective synthetic lethal target in BRG1-deficient cancer, and vice-versa (101,102). Despite the compensatory nature of BRG1/BRM in many tumorigenic contexts, concomitant loss of expression of the ATPases has been described in metastatic murine models and patient-derived non-small-cell lung cancer (NSCLC) cell lines and is associated with poor patient survival (91,103,104). In *C elegans*, the sole SWI/SNF ATPase, *swsn-4,* has a high degree of homology to both mammalian BRG1 and BRM, providing a unique opportunity to accessibly model the connection between the dual loss of BRG1/BRM associated with poor prognostic outcomes in NSCLC and cellular invasion in the AC. Additionally, a recent study demonstrated a phenotypic dosage-sensitivity following loss of SWI/SNF expression in relation to terminal differentiation of *C. elegans* muscle and vulval tissues (35), suggesting that the context- and dose-dependent relationship seen in mammalian development and SWI/SNF-deficient cancers may represent a more general behavior of the complex.

Here we used the *C. elegans* AC invasion system as a model to investigate whether the dose-dependent relationship between the ATPase and differentiated phenotype extends to cellular invasion. The first indication that the functional dose of SWI/SNF may have an instructive role in AC invasion was the relative enhancement of all endogenously tagged subunits of the complex in the AC relative to neighboring VU and VPC tissues (**Fig S4**). While it is tempting to interpret the enhancement of SWI/SNF subunit expression in the AC as evidence for the dependence of cellular invasion on SWI/SNF activity, it is also possible that this difference in expression is a consequence of terminal differentiation, since at the time of invasion, the AC is terminally differentiated unlike the VU and VPC). Further investigation is required to determine whether endogenously tagged SWI/SNF subunits are enhanced in the AC relative other terminally differentiated cells.

Additionally, we find that some degree of SWI/SNF intra- and inter-complex regulation exists in the AC at the time of invasion with both SWI/SNF assemblies cooperating to activate expression of the ATPase (**Fig S5 B**). It is possible that this added level of complex autoregulation contributes to an “optimal” dose of the ATPase in a cell- and context-specific manner. It is important to note that the model proposed in Fig S5 F is incomplete and is included to describe the translational consequence of individual SWI/SNF subunits after transcriptional knockdown of other individual SWI/SNF subunits. However, these data can be meaningfully used to inform interpretations of results for our enhanced SWI/SNF RNAi experiments in the AC.

To directly correlate SWI/SNF dose and invasive phenotype in the AC, we used a combination of an enhanced *swsn-4(RNAi)* vector with a tissue-specific antiGFP-targeting nanobody construct. These combined technologies allowed us to titrate the dose of ATPase in the AC at the time of invasion. By assessing AC invasion phenotypes at wildtype levels of SWSN-4 and in moderate and severe ATPase knockdown conditions, results indicate that cellular invasion and cell cycle control depends on the dose of functional SWI/SNF present in the AC.

In addition to reflecting the dose-dependent nature of the SWI/SNF ATPase in cancer, our data in the AC is consistent with work done in *C. elegans* early mesoblast development where complete loss of the *swsn-4* ATPase using a catalytically dead mutant and lineage-specific knockout strategy results in loss of cell cycle arrest (35). Although we cannot be sure that combining *swsn-4(RNAi)* with an antiGFP-targeting nanobody to deplete the SWI/SNF ATPase results in complete loss of protein expression, we show that treatment with the improved *swsn-4(RNAi)* vector alone is sufficient to phenocopy the null phenotype previously reported in late mesoblast (SM) development (**Fig S6**). Additionally, enhancement of the AC mitotic phenotype statistically tracked with a progressive step down in mean expression of the ATPase in the AC across our experiments. Using endogenous fluorescent reporters for conserved pro-invasive genes, we also provide evidence that the SWI/SNF complex is involved broadly with the regulation of the cell cycle-dependent and -independent subcircuits of the AC GRN (**Fig 6**). Altogether, this data supports the hypothesis that SWI/SNF cell-autonomously contributes to cell cycle control in a dose-dependent manner and provides the first line of evidence to link SWI/SNF ATPase dosage to the dichotomy between invasion and proliferation (**Fig 8**).

**Figure 8.**
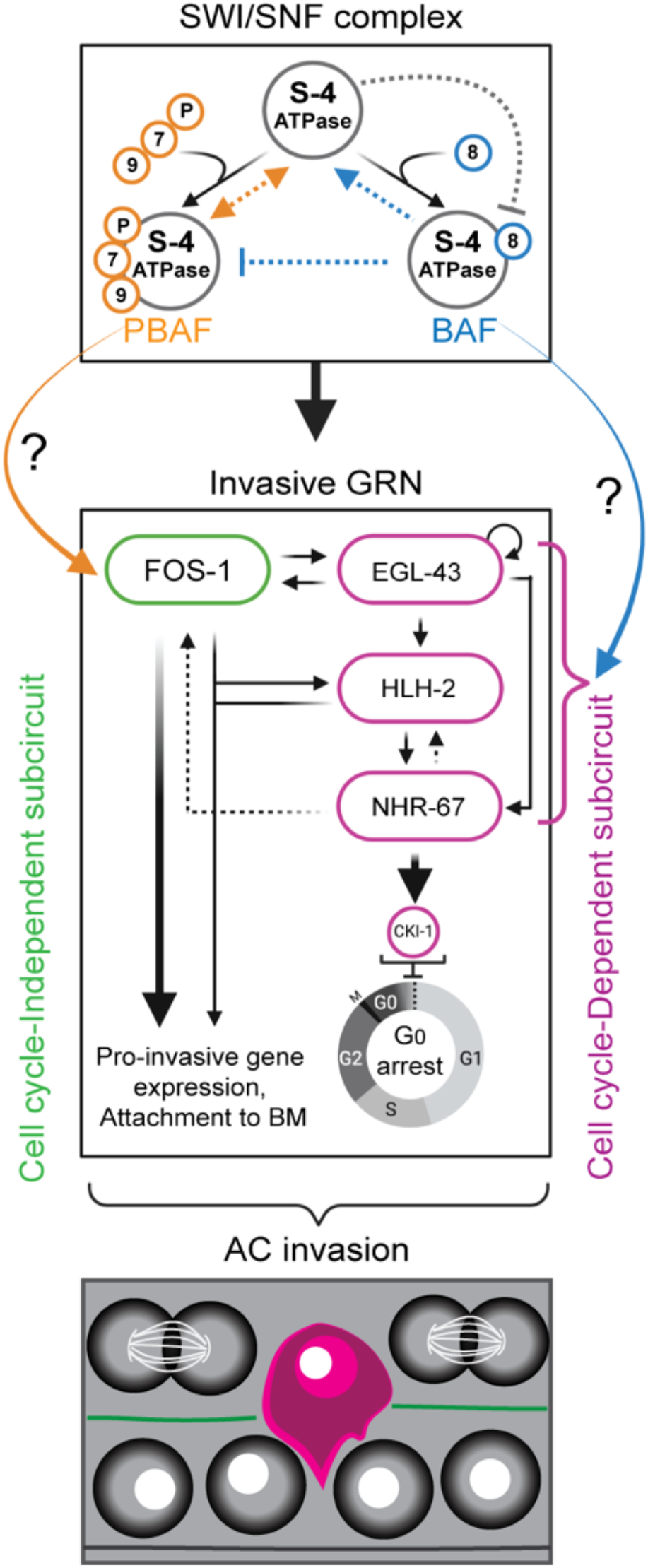
SWI/SNF complex promotes AC invasion. Schematic summary of the how the SWI/SNF ATPase (S-4*, swsn-4*), PBAF (orange – S-7, swsn*-7*; S-9, *swsn-9*; P, *pbrm-1*), and BAF (blue – S-8, *swsn-8*) assemblies contribute to AC invasion.

### The SWI/SNF BAF assembly promotes cellular invasion through induction of G_0_ cell cycle arrest

While previous work in our lab, based on localization of a DNA licensing factor, CDT-1, has demonstrated indirectly that ACs must arrest in a G_0_/G_1_ cell cycle state (22,24), we lacked a sensitive enough tool to distinguish between these two interphase states. From our recent work utilizing a CDK sensor to examine the proliferation-quiescence decision in *C. elegans*, we can distinguish between pre-terminal cells in the somatic gonad in G_1_ (mean C/N ratio: 0.67+/-0.10) as compared to terminally differentiated G_0_ uterine cells (mean C/N ratio: 0.30+/-0.11) (36). Here, we compare CDK activity measurements in the ACs of control animals with that of the terminal Pi lineage to provide the first quantitative demonstration that ACs arrest in a CDK^low^ G_0_ state to invade (**Fig 4 A, E, F**). Furthermore, by combining the CDK sensor with loss of SWI/SNF subunits, our data indicate that the SWI/SNF BAF assembly is specifically responsible for regulation of G_0_ cell cycle arrest in the AC.

The SWI/SNF core complex is composed of an ATPase, core, and accessory factors, which are thought to collectively provide a platform of common factors that is bound by assembly-specific subunits in a mutually exclusive manner. In *C. elegans*, the association of SWSN-8 (BAF250a/ BAF250b/ ARID1A/ ARID1B) with the common factors forms the SWI/SNF BAF assembly. Since we identified representative subunits from the core of the complex and both SWI/SNF BAF and PBAF assemblies in our CRF RNAi screen, we took a cell biological and genetic approach to investigate the role of the each SWI/SNF assembly in promoting cellular invasion. Here, using a DHB-based CDK sensor (36,70), we show that loss of either core or BAF assembly subunits specifically results in mitotic ACs that failed to invade the BM. Our cell cycle sensor data first establishes that wild-type AC invades in a G_0_ CDK^low^ state, and second, that a major contribution of the BAF assembly to AC invasion is through maintenance of this G_0_ arrest, as many ACs that failed to invade the BM had increasing CDK activity, indicative of cells cycling in G_1_, S or G_2_. Alternatively, 14% of ACs that failed to invade the BM following loss of *swsn-4*/ATPase of the complex had CDK activity ratios indicative of G_0_ cell cycle arrest, suggesting a cell cycle-independent defect. In support of this, forced arrest of BAF-deficient ACs in G_0_ was sufficient to significantly rescue invasion, whereas CKI-1 induction failed to rescue invasion in ACs with RNAi-mediated loss of *swsn-4*/ATPase. Altogether, our results indicate that the SWI/SNF complex contributes to AC invasion through regulation of G_0_ cell cycle arrest via the BAF assembly. Further investigation will require biochemical techniques to identify cell cycle regulators and TF targets of the BAF assembly to provide a mechanistic explanation for how exactly BAF regulates the chromatin landscape to promote invasion (**Fig 8**, blue arrow). Targeted DNA adenine methyltransferase identification (TaDa) is an attractive biochemical approach that may be adaptable to the AC invasion system, as this approach has been characterized as an effective, tissue-specific method to identify TF-target sequence interactions in the *C. elegans* epidermis (106).

### The PBAF assembly regulates AC invasion and attachment to the BM

Based on homology and phenotypic characterization in this study and previous publications, the *C. elegans* PBAF assembly consists of the PBRM-1 (PBRM1/BAF180), SWSN-7 (BAF200/ARID2), and SWSN-9 (BRD7/BRD9) in association with the SWI/SNF common factors. Previous work in *C. elegans* has not revealed a connection between the PBAF assembly and cell cycle arrest. Our initial experiments with improved RNAi vectors targeting PBAF subunits resulted in a lower penetrance of AC invasion defects relative to loss of core or BAF subunits. Additionally, our CDK sensor data suggested that non-invasive ACs deficient in *pbrm-1* remain in a G_0_ cell cycle state. Thus, our data shows no PBAF contribution to cell cycle control in the AC. To confirm this, we used the auxin inducible degron (AID) system to robustly deplete the PBAF assembly through combined loss of endogenous PBRM-1::mNG::AID with RNAi-mediated knockdown of either of the other two PBAF assembly subunits, *swsn-7* or *swsn-9*. This combination knockdown strategy corroborated our previous results as we saw no significant penetrance of extra ACs. Rather, here we associate a striking AC detachment phenotype with strong combined knockdown of the PBAF assembly subunits. We also note aberrant BM morphology in some ACs deficient in PBAF subunits, with only one of the two BMs removed, suggesting that this assembly regulates attachment and extracellular matrix (ECM) remodeling in wild-type ACs to promote invasion. We hypothesize that the PBAF assembly is regulating ventral BM attachment and ECM remodeling potentially through the regulation of HIM-4/Hemicentin, an extracellular immunoglobulin-like matrix protein that functions in the AC to fuse the two BMs through the formation of a novel BM-BM adhesion, the B-LINK (107). Finally, although RNAi-mediated transcriptional knockdown of PBAF assembly subunits only partially depleted levels of FOS-1::GFP, a key TF responsible for the expression of MMPs and other pro-invasive targets, we detected significant enhancement of invasion defects when depleting *fos-1* in a putative hypomorphic *pbrm-1* background. Reciprocally, depletion of *pbrm-1* enhanced the invasion defect of a quintuple MMP mutant. Since we noted multiple instances of AC-BM detachment following PBAF assembly subunit depletion, we propose that PBAF functions in part with FOS-1 to facilitating activating chromatin states at the regulatory regions of pro-invasive genes required for BM attachment. Future studies should include an investigation of PBAF assembly interactions with other FOS effectors and interactors at the transcriptional level. Additionally, given the broad genomic regulation exhibited by chromatin remodelers, biochemical techniques such as chromatin immunoprecipitation (ChIP) and single cell RNA sequencing will be crucial going forward to generate a comprehensive roster of direct and indirect targets of each SWI/SNF assembly in the AC. Collectively, these results reveal a distinct contribution for each SWI/SNF assembly to the process of cellular invasion at the phenotypic level and reiterate the dependence of each assembly on the functional dose of ATPase in a cell.

## ACKNOWLEDGEMENTS

We are thankful to David Gray, Ed Luk, Laura Mathies, Benjamin Martin, Robert Morabito, Valerie Reinke, Courtney Tello, and Gerald Thomsen for advice and comments on this manuscript. We would also like to thank Thom Geer of Nobska Imaging for advice and ‘scientific enabling’. Some *C. elegans* strains were provided by the CGC, which is funded by NIH Office of Research Infrastructure Programs (P40 OD010440).

## AUTHOR CONTRIBUTIONS

J.J.S. and D.Q.M. designed the experiments. J.J.S., Y.X., M.A.Q.M., and T.N.M.-K. performed the experiments. J.J.S., A.Q.K., and M.C. organized the CRF RNAi screen. J.J.S., M.C., N.P., and Y.X. performed the RNAi screen. J.J.S., A.Q.K., S.L., T.N.M.-K., Y.X., K.W. and P.K generated strains. J.J.S performed the data analysis and prepared the manuscript with feedback from other authors. J.J.S. and D.Q.M acquired funding for the completion of this project.

## DECLARATION OF INTERESTS

The authors declare no competing interests.

## FUNDING

This work was funded by the National Institute of General Medical Sciences (NIGMS) [1R01GM121597-01 to D.Q.M.]. D.Q.M. is also a Damon Runyon-Rachleff Innovator supported (in part) by the Damon Runyon Cancer Research Foundation [DRR-47-17]. J.J.S was supported by an NIH/NIGMS Diversity Supplement [3R01GM121597-02S1] and the W. Burghardt Turner Fellowship. M.A.Q.M. [3R01GM121597-03S1] and F.E.Q.M. [3R01GM121597-04S1] are supported by an NIH/NIGMS Diversity Supplement. A.Q.K. [F31GM128319], R.C.A [1F31GM1283190], and T.N.M.-K. [F31HD100091-01] were supported by the NIH. N.J.P [132969-PF-18-226-01-CSM] was supported by the American Cancer Society (ACS). P.K. [R01NS118078] is supported by the National Institute of Neurological Disorders and Stroke.

## MATERIALS & METHODS

### *C. elegans* strains and culture conditions

All animals were maintained under standard conditions and cultured at 25°C, except strains containing temperature-sensitive alleles *swsn-1(os22), swsn-4(os13)*, and the uterine-specific RNAi hypersensitive strain used in the chromatin remodeler screen containing the *rrf-3(pk1426)* allele, which were maintained at either 15°C or 20°C (108). The heat shock inducible *cki-1::mTagBFP2* transgene was expressed via incubating animals at 32°C for 2-3 hours in a water bath starting at the P6.p 2-cell VPC stage. Animals were synchronized for experiments through alkaline hypochlorite treatment of gravid adults to isolate eggs (109). In the text and figures, we designate linkage to a promoter through the use of a (p) and fusion of a proteins via a (::) annotation.

### Molecular biology and microinjection

SWI/SNF subunits *swsn-4* and *swsn-8* were tagged at their endogenous loci using CRISPR/Cas9 genome editing via microinjection into the early adult hermaphrodite syncytial gonad (110,111). Repair templates were generated as synthetic DNAs from either Integrated DNA Technologies (IDT) as gene blocks (gBlocks) or Twist Biosciences as DNA fragments and cloned into *ccdB* compatible sites in pDD282 by New England Biolabs Gibson assembly (112). Homology arms ranged from 690-1200 bp (see **Tables S5** for additional details). sgRNAs were constructed by EcoRV and NheI digestion of the plasmid pDD122. A 230 bp amplicon was generated replacing the sgRNA targeting sequence from pDD122 with a new sgRNA and NEB Gibson assembly was used to generate new sgRNA plasmids (see **Table S5** for additional details). Hermaphrodite adults were co-injected with guide plasmid (50 ng/μL), repair plasmid (50 ng/μL), and an extrachromosomal array marker (pCFJ90, 2.5 ng/μL), and incubated at 25 °C for several days before screening and floxing protocols associated with the SEC system (112).

### RNA interference (RNAi)

All 269 RNAi clones assessed in the chromatin remodeler screen were derived from the commercially available Vidal or Ahringer RNAi libraries. Presence of inserts into the L4440 RNAi vector was confirmed via colony PCR amplification of all L4440 vectors used in the chromatin remodeler screen. Vectors which resulted in penetrant loss of invasion (see **Table S2**) were also sequenced to confirm the identity of the insert targeting chromatin remodeler genes in the L4440 vector using Sanger sequencing at the Genomics Core Facility at Stony Brook University. An RNAi sub-library of SWI/SNF subunits was constructed by cloning 950-1000 bp of synthetic DNA based on cDNA sequences available on WormBase (www.wormbase.org) into the highly efficient T444T RNAi vector (113,114). Synthetic DNAs were generated by Twist Biosciences as DNA fragments and cloned into restriction digested T444T using NEB Gibson Assembly (see **Tables S6** for additional details). For all experiments, synchronized L1 stage animals were directly exposed to RNAi through feeding with bacteria expressing dsRNA (115).

### Auxin-mediated degradation

To combine RNAi with the depletion of AID-tagged proteins, 1 mM K-NAA was used, and its effects were analyzed as previously described(116). Briefly, L1 animals were first synchronized by sodium hypochlorite treatment and transferred to NGM plates seeded with the RNAi vector of interest. At the P6.p 1-cell stage, a time in development where the AC has already undergone specification, animals were transferred to RNAi-seeded plates treated with K-NAA. Animals were staged by DIC.

### Live cell microscopy

All micrographs included in this manuscript were collected on a Hamamatsu Orca EM-CCD camera mounted on an upright Zeiss AxioImager A2 with a Borealis-modified CSU10 Yokagawa spinning disk scan head using 405nm, 488 nm, and 561 nm Vortran lasers in a VersaLase merge and a Plan-Apochromat 100x/1.4 (NA) Oil DIC objective. MetaMorph software (Molecular Devices) was used for microscopy automation. Several experiments and all RNAi screening were scored using epifluorescence visualized on a Zeiss Axiocam MRM camera, also mounted on an upright Zeiss AxioImager A2 and a Plan-Apochromat 100x/1.4 (NA) Oil DIC objective. Animals were mounted into a drop of M9 on a 5% Noble agar pad containing approximately 10 mM sodium azide anesthetic and topped with a coverslip.

### Assessment of AC invasion

Both for the purposes of the CRF RNAi screen and all other experiments AC invasion was scored at the P6.p 4-cell stage, when 100% of wild-type animals exhibit a breach in the BM (14). In strains with the laminin::GFP transgene, an intact green fluorescent barrier under the AC was used to assess invasion. Wild-type invasion is defined as a breach as wide as the basolateral surface of the AC (14). Raw scoring data is available in **Tables S1 and S4.**

### Image quantification and statistical analyses

Images were processed using Fiji/ImageJ (v.2.1.0/1.53c) (117). Expression levels of GFP::SWSN-4, SWSN-8::GFP, PBRM-1::eGFP, and PBRM-1::mNG::AID were measured by quantifying the mean gray value of AC nuclei, defined as somatic gonad cells near the primary vulva expressing the *cdh-3p::mCherry::moeABD* transgene. Background subtraction was performed by rolling ball background subtraction (size=50). For characterization of experiments involving SWI/SNF endogenous tags and AC GRN TFs::GFP treated with *SWI/SNF(RNAi)* and GFP-targeting nanobody the L3 stage, only animals exhibiting defects in invasion were included in the analysis. Data was normalized to negative control (empty vector) values for the plots in **Fig 3** and **Fig S8**. Quantification of either CDK cell cycle sensor (either DHB::GFP or DHB::2xmKate2) was performed by hand, as previously described (36). Images were overlaid and figures were assembled using Adobe Photoshop 2020 (v. 21.1.2) and Adobe Illustrator 2020 (v. 24.1.2), respectively. Statistical analyses and plotting of data were conducted using RStudio (v. 1.2.1335). Statistical significance was determined using either a two-tailed Student’s t-test or Fisher’s exact probability test. Figure legends specify when each test was used and the p-value cut-off.

## SUPPLEMENTAL FIGURES

**Figure S1.**
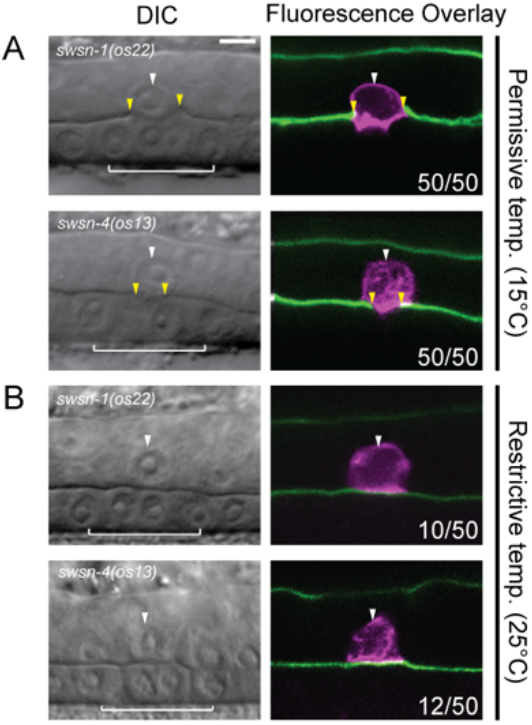
AC invasion is disrupted in temperature sensitive SWI/SNF hypomorphs. Single planes of confocal z-stacks representing AC invasion in *swsn-1(os22) and swsn-4(os13)* temperature sensitive mutants with fluorescently labeled AC (magenta, *cdh-3>mCherry::moeABD*) and BM (green, *laminin::GFP*) scored at the permissive temperature **(A)** and restrictive temperature **(B)**. Significant loss of invasion was seen in both *swsn-1(os22)* (20% loss of invasion) and *swsn-4(os13)* (24% loss of invasion) hypomorphic^ts^ strains when grown at the restrictive temperature 25°C and assessed at the P6.p 4-cell 1° VPC stage **(B)**.

**Figure S2.**
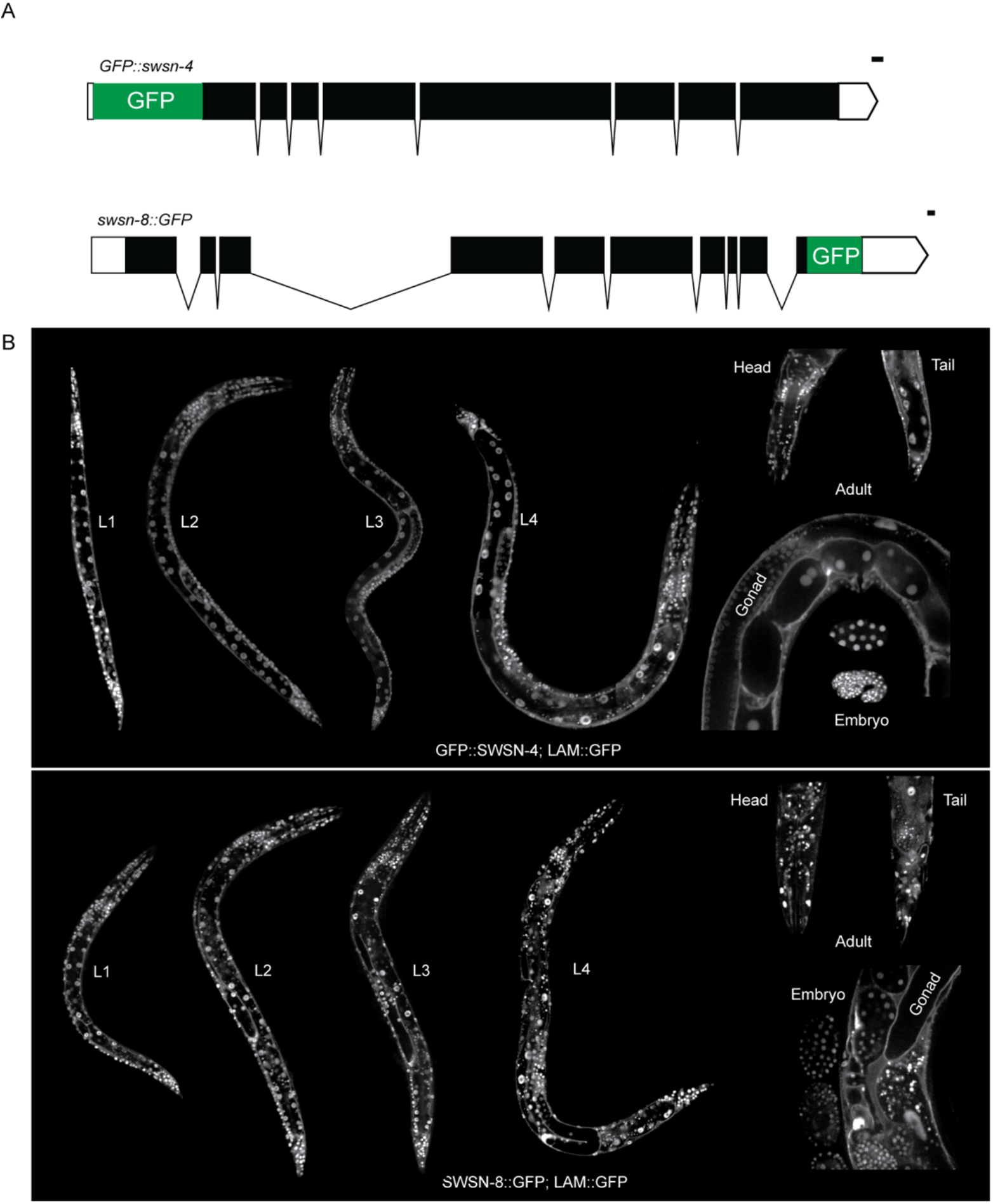
Endogenous SWSN-4::GFP and SWSN-8::GFP are ubiquitously expressed. **(A)** Schematic (from http://wormweb.org/exonintron) depicting GFP insertion into the endogenous N and C termini of *swsn-4* (top) and *swsn-8* (bottom), respectively. Scale bar, 100 bp. **(B)** Expression of endogenously GFP-labeled SWI/SNF ATPase SWSN-4 protein and BM (*laminin::GFP*) in all larval stages (L1-L4), adult animals, and embryos. **(C)** Expression of endogenously GFP-labeled SWI/SNF ATPase SWSN-4 protein and BM (*laminin::GFP*) in all larval stages (L1-L4), adult animals, and embryos. Images in B-C are not to scale.

**Figure S3.**
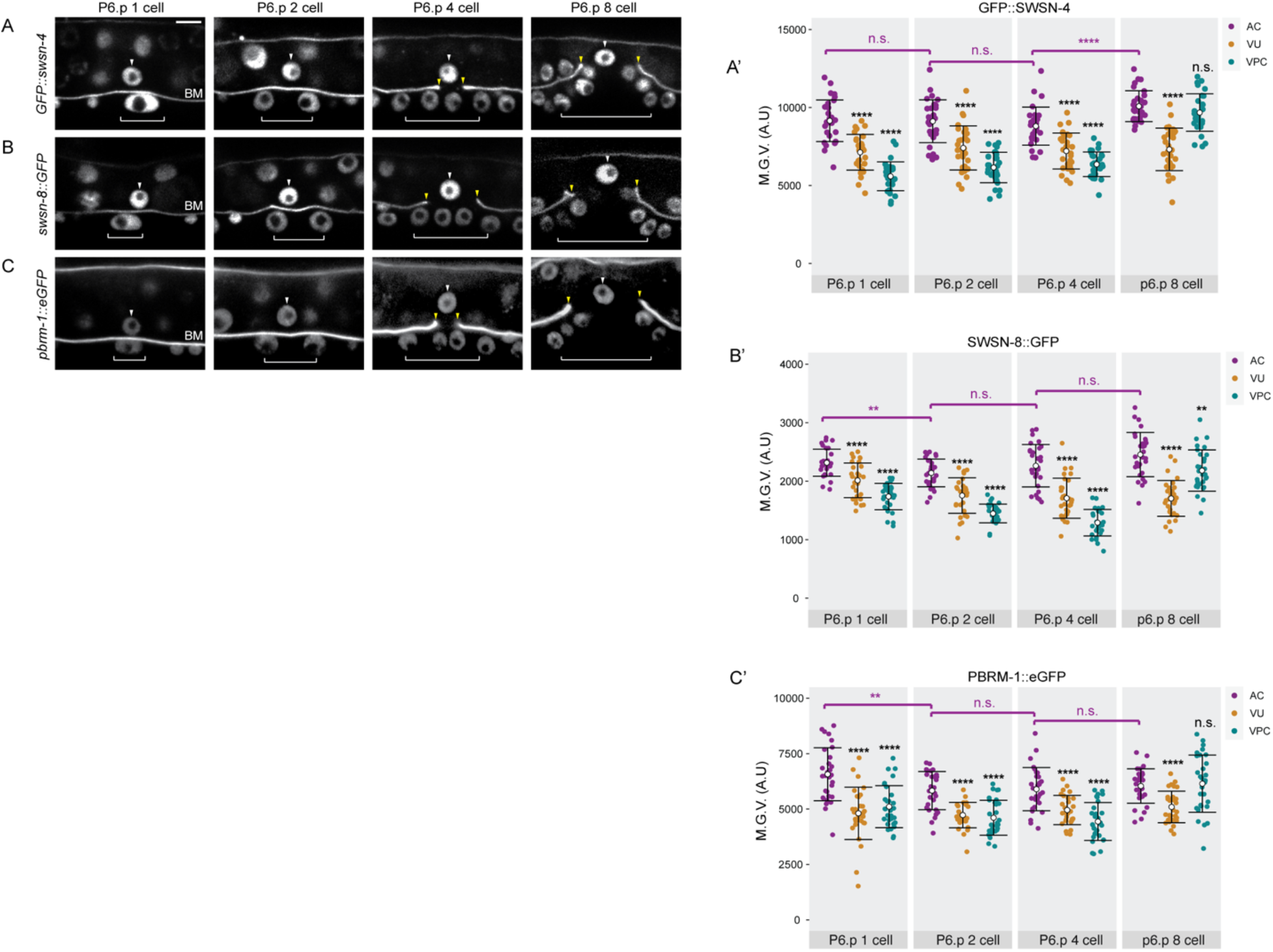
Fluorescent knock-ins express in the AC pre-, during, and post-invasion. Fluorescent micrographs depicting expression of SWSN-4::GFP **(A),** SWSN-8::**GFP (B)**, and PBRM-1::eGFP **(C)** in the AC, VU, and VPCs from the P6.p 1 cell to the P6.p 8 cell stages of development and corresponding quantifications. White arrowheads indicate AC, White brackets indicate 1° VPC stage. **(A’-C’)** Quantification of endogenous GFP expression of SWI/SNF subunit in the AC, VU, and VPC over time. Statistical comparisons were made between the expression of each SWI/SNF subunit in the AC over time (magenta bracket and asterisks) or between the expression of each subunit in the AC relative to the expression of the same subunit in the neighboring VPCs or VUs at the same time (black asterisks) using Student’s *t*-test (n≥30 for each stage and subunit; p values are displayed above compared groups). n.s. not significant.

**Figure S4.**
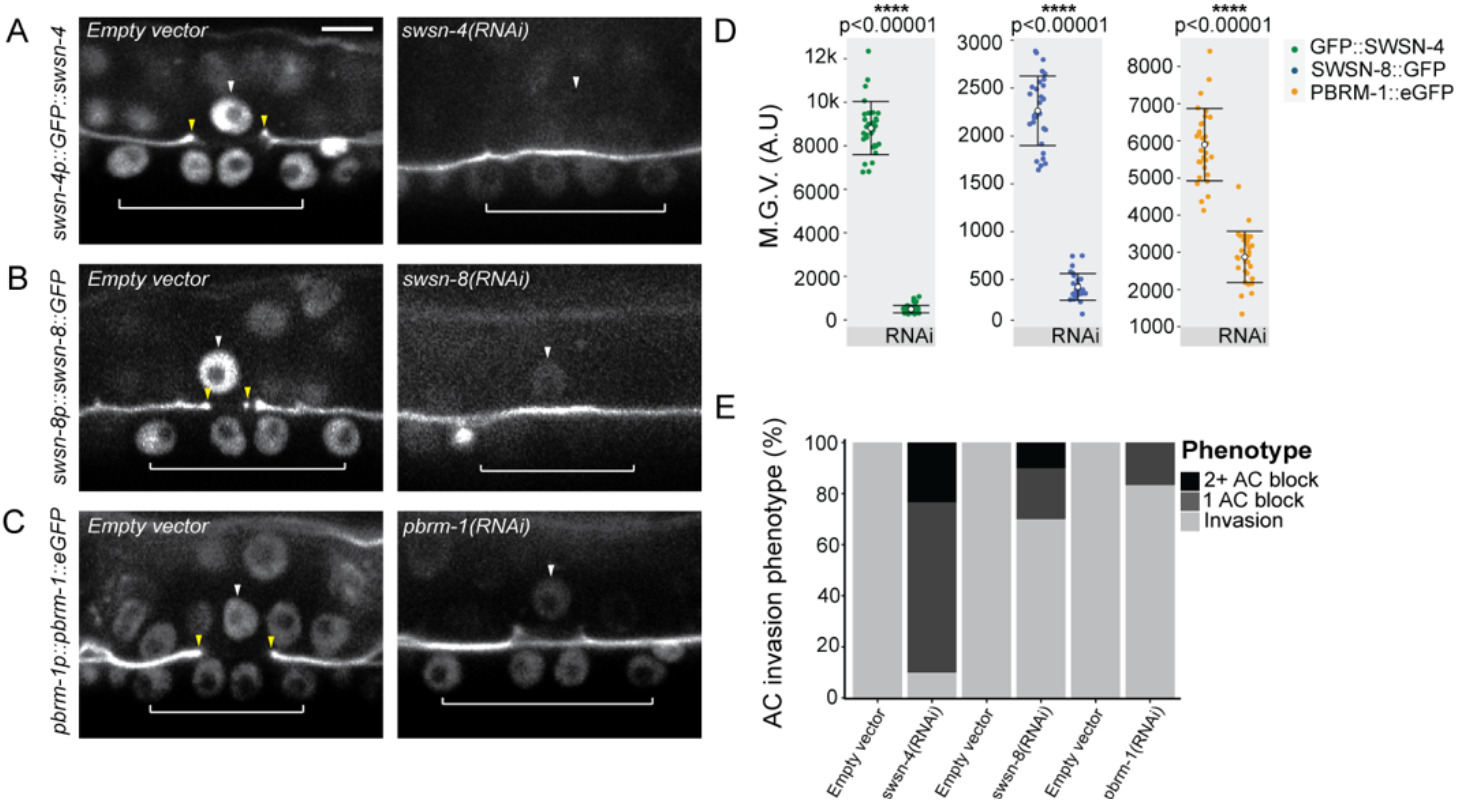
Improved SWI/SNF RNAi significantly knocks down SWI/SNF expression in the AC. Fluorescent micrographs depicting BM (*laminin::GFP*) and expression of SWSN-4::GFP **(A)**, SWSN-8::GFP **(B)**, and PBRM-1::eGFP **(C)** in the AC in animals fed empty vector control (left) or RNAi targeting the endogenous allele (right). White arrowheads indicate AC(s), yellow arrowheads indicate boundaries of breach in BM, and white brackets indicate 1 VPCs. **(D)** Corresponding quantifications of fluorescent expression. Statistical comparisons were made between the expression of each SWI/SNF subunit in the AC in control and RNAi-treated animals using Student’s *t*-test (n≥30 for each stage and subunit; p values are displayed above compared data). **(E)** Stacked bar chart showing percentage of AC invasion defects corresponding to each treatment, binned by AC phenotype (n≥30 animals per condition).

**Figure S5.**
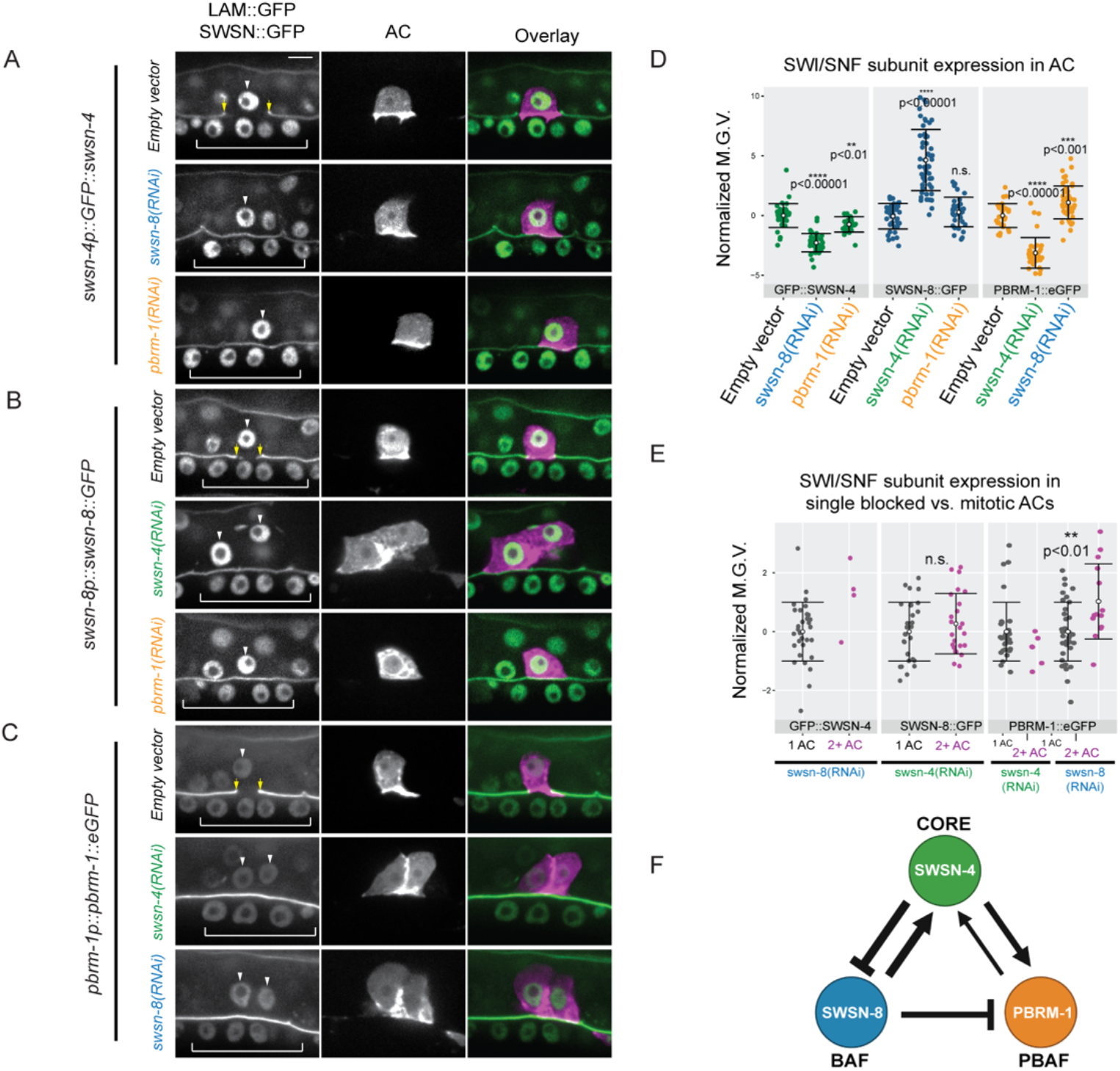
SWI/SNF subunits exhibit intra-complex and inter-assembly regulation. **(A-C)** Representative fluorescence micrographs depicting endogenous GFP expression of individual SWI/SNF subunits representative of the core (*swsn-4*, **A**), BAF assembly (*swsn-8,* **B**), and PBAF assembly (*pbrm-1*, **C**) in the AC (*cdh-3p::mCherry::*moeABD) following treatment with targeting either SWI/SNF assembly **(A),** or the core ATPase and alternative SWI/SNF assembly **(B-C)**. White arrowheads indicate AC(s), yellow arrowheads indicate boundaries of breach in BM, and white brackets indicate 1 VPCs. **(D)** Quantification of fluorescence expression (mean gray value) of endogenous subunits in each condition. Statistical comparisons were made between the expression of each SWI/SNF subunit in the AC in control and RNAi-treated animals using Student’s *t*-test (n≥30 for each stage and subunit; p values are displayed above compared data). n.s. not significant**. (E)** Quantification of fluorescence expression of endogenous GFP-tagged subunits of non-invasive ACs following loss of expression of alternative SWI/SNF subunits, binned per RNAi treatment by phenotype into single non-invasive AC (1AC) and mitotic non-invasive AC (2+ AC). Statistical comparisons (Student’s *t*-test; p values are displayed above compared data) were limited to conditions with n>10 ACs in each phenotype. n.s. not significant. **(F)** Schematic summary of SWI/SNF core and assembly auto and cross regulation.

**Figure S6.**
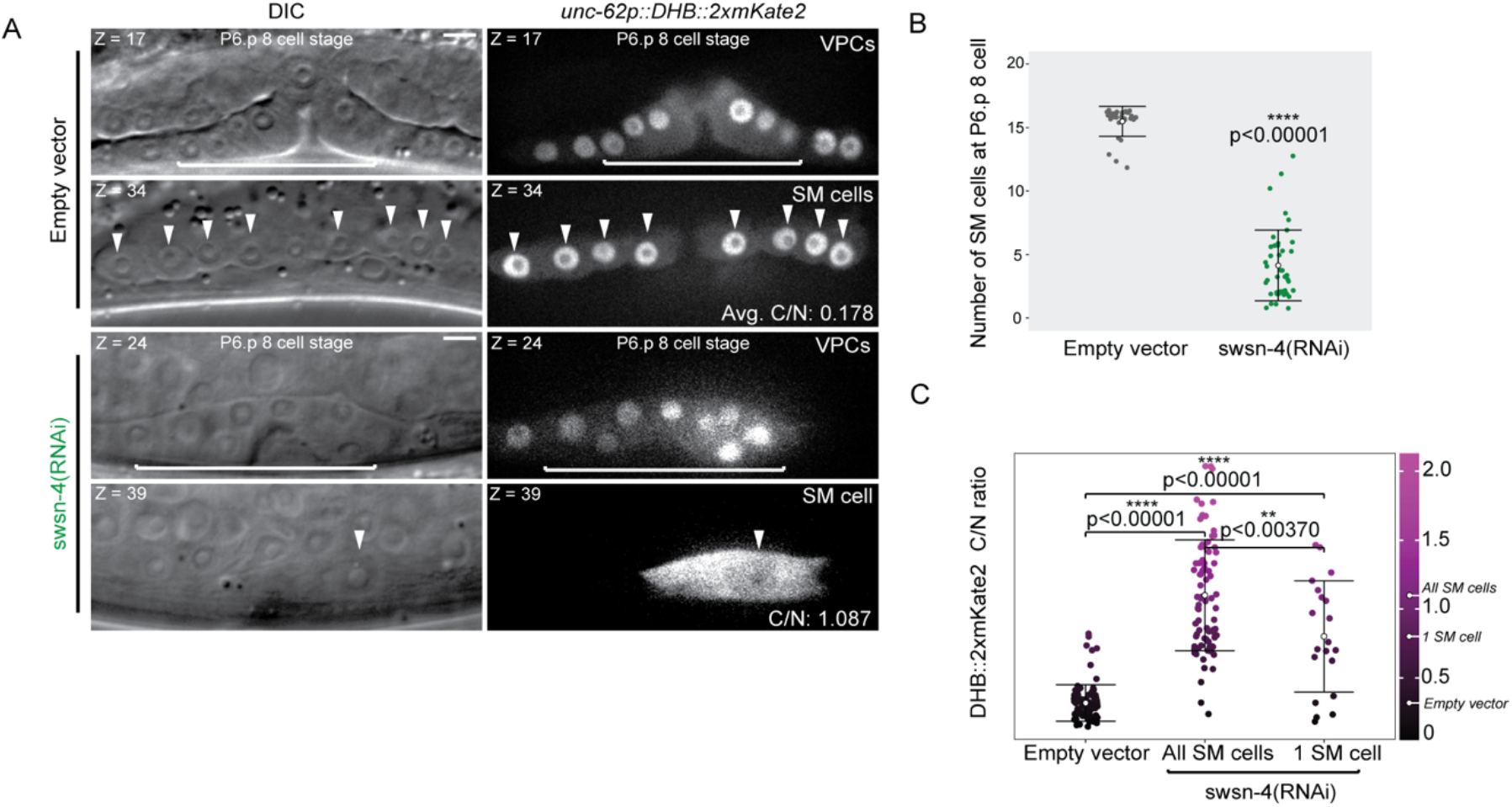
Improved *swsn-4* RNAi recapitulates SWI/SNF ATPase null phenotype in the sex myoblasts. **(A)** Single confocal z-planes depicting DIC (left) and expression of lineage-restricted CDK sensor (*unc-62>DHB::2xmKate2*, right) in the vulva and SM cells at the P6.p 8 cell stage corresponding to the stage when wild-type SM cells differentiate and exit the cell cycle. Animals were treated with empty vector control (top) or *swsn-4(RNAi)* (bottom). All representative images in each treatment are derived from the same z-stack from the same animal in the corresponding z-plane (top-left). Average or individual C/N CDK sensor ratios are listed in the bottom-right of corresponding panels. White arrowheads indicate individual SM cells. White brackets indicate 1° VPCs. **(B)** Quantification of the number of SM cells present at the P6.p 8 cell stage in control (black) and *swsn-4(RNAi)* treated animals (SMs arrive on both left and right side of vulva at the P6.p 8 cell, green; SMs arrive on either left or right side of vulva at the P6.p 8 cell stage, blue). **(C)** C/N CDK sensor ratios for SM cells in each treatment. Mean C/N ratio is represented by colored open circles and correspond to numbers of the same color. Gradient scale depicts cell cycle state as determined by quantification of each AC in all treatments (n≥30 animals per treatment), with dark-black depicting differentiation into G_0_/G_1_ and lighter-magenta depicting G_2_ cell cycle states.

**Figure S7.**
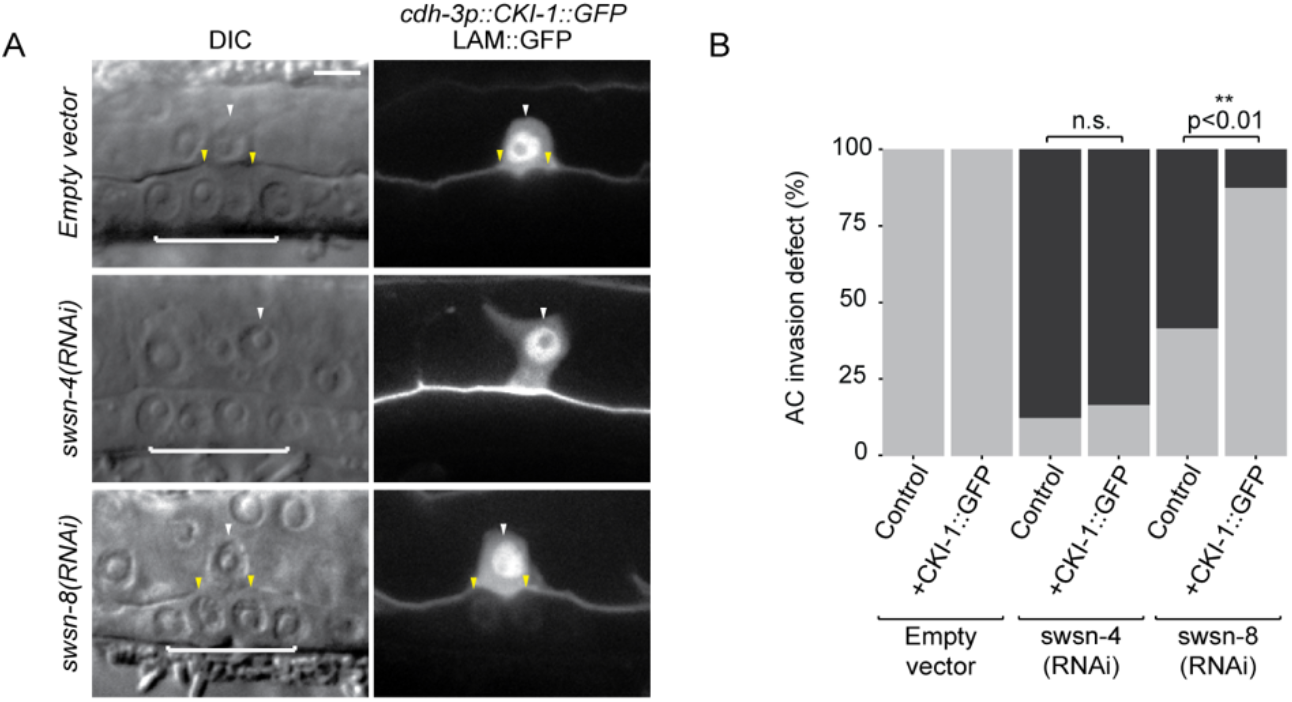
AC-specific expression of CKI-1 rescues invasion in BAF-depleted ACs. **(A)** DIC (left) and fluorescent (right) images depicting BM (*laminin::GFP*) and AC-specific CKI-1 (*cdh-3>CKI-1::GFP*) in empty vector control animals (top) and animals treated with *swsn-4(RNAi)* (middle) or *swsn-8(RNAi)* (bottom). **(B)** Stacked bar chart showing quantification of percentage of AC invasion defects corresponding to each treatment (n≥30 animals per condition, p values for Fisher’s exact test comparing invasion penetrance in control animals and animals with the rescue transgene (+CKI-1::GFP) are displayed above black brackets).

**Figure S8.**
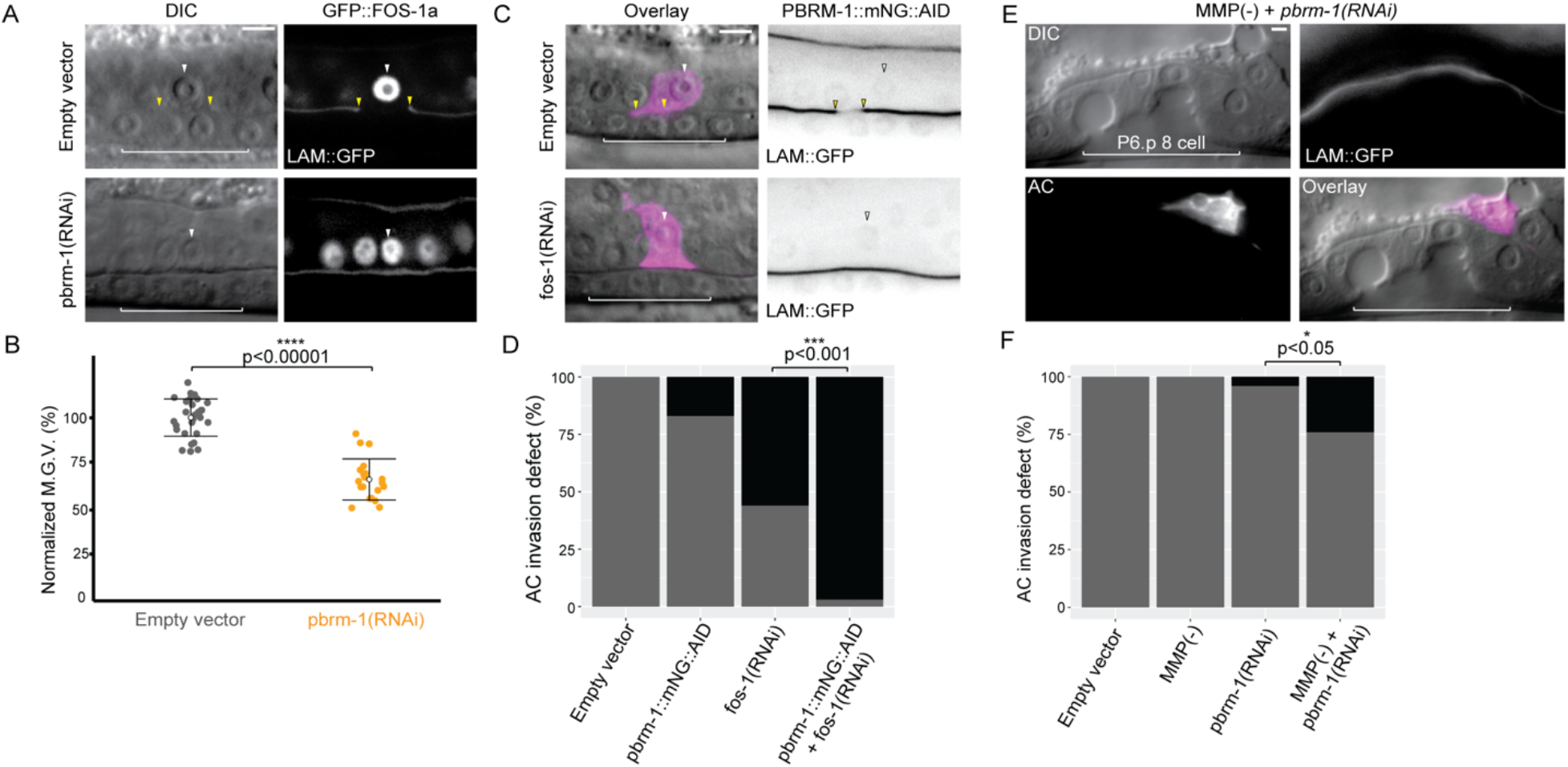
PBAF partially regulates the FOS-1 transcription factor. **(A)** Representative DIC (left) and fluorescent (right) micrographs depicting expression of endogenous GFP::FOS-1a and BM (*laminin*:*:GFP*) in control (top) and *pbrm-1(RNAi)* treated (bottom) animals. **(B)** Quantification of GFP::FOS-1a expression in ACs of control and *pbrm-1(RNAi)* treated animals, normalized to mean expression of control group. Statistical comparisons were made between expression in the AC in control and RNAi-treated animals using Student’s *t*-test (n≥20 for each condition; p value is displayed above black bracket). **(C)** DIC-Fluorescence overlay (left), and PBRM-1::mNG::AID and BM (LAM::GFP) (right), in animals treated with empty vector control (top) or *fos-1(RNAi)* (bottom). **(D)** Stacked bar chart showing percentage of AC invasion defects corresponding to each treatment and genetic background in C (n≥30 animals per condition, p values for Fisher’s exact test comparing invasion defect penetrance in wild-type animals treated with *fos-1(RNAi)* and *pbrm-1::mNG::AID* animals treated with *fos-1(RNAi)* is displayed above black bracket). **(E)** Representative DIC (top-left), BM (LAM::GFP, top-right), AC (*cdh-3>PH*, bottom-left), and overlay (bottom-right) of P6.p 8 cell vulva in an MMP-deficient (-) animal treated with *pbrm-1(RNAi)*. **(F)** Stacked bar chart showing percentage of AC invasion defects corresponding to each treatment and genetic background in E (n≥30 animals per condition, p values for Fisher’s exact test comparing invasion defect penetrance in wild-type animals treated with *pbrm-1(RNAi)* and MMP(-) animals treated with *pbrm-1(RNAi)* is displayed above black bracket).

## SUPPLEMENTAL TABLES

**Table S1. CRFs assessed for AC invasion contribution** (see excel file)

270 chromatin regulating factors targeted by RNAi for AC invasion defects. n≥30 animals for each RNAi clone. For each RNAi clone tested, the corresponding genetic sequence name, public name, protein annotation, and human homolog (HUGO Gene Nomenclature) from www.wormbase.com is given. Penetrance for each invasion defects is given as the % of animals with ACs that fail to invade the BM at the P6.p 4 cell stage out of the total number of animals assessed (Block/Invasion+Partial). Partial refers to cases where an animal had a breach in the BM narrower than the width of the basolateral surface of the invading AC. Genes in bold were recovered as significant regulators of AC invasion (Table S2). Annotations were mined from the STRING consortium www.string-db.org. Asterisks in human ortholog column denote genes with > 5 detected human orthologs, for which only the first 5 returned orthologs were listed. N.A. denotes genes for which no human ortholog exists. List is organized alphabetically based on genetic sequence name.

**Table S2. Significant regulators of AC invasion** (see excel file)

41 chromatin and chromatin regulating factors (CRFs) identified as significant regulators of AC invasion. For each RNAi clone listed, the corresponding genetic sequence name, public name, and human homolog is listed. AC invasion scoring data is provided for each clone at the P6.p 4 cell stage. Genes were determined to be significant AC invasion regulators if RNAi targeting resulted in ≥ 20% loss of invasion at the P6.p 4-cell stage (n≥30 animals). Genes in bold are components of the SWI/SNF complex. Asterisks denote genes previously published to regulate *C. elegans* AC invasion. N.A. denotes genes for which no human ortholog exists. List is organized alphabetically based on genetic sequence name.

**Table S3. Enhanced (T444T) RNAi vectors used in this study** (see excel file)

**Table S4. Strains used in this study** (see excel file)

**Table S5. CRISPR reagents** (see excel file)

## Notes

### Competing Interest Statement

The authors have declared no competing interest.

### Summary of Updates

updated version with supplemental tables

